# Lineage tracing via associative chromosome/plasmid barcoding with siBar

**DOI:** 10.1101/2022.03.17.484700

**Authors:** Han Mei, Anton Nekrutenko

## Abstract

We describe a new method, siBar, which enables simultaneous tracking of plasmids and chromosomal lineages within a bacterial host. siBar involves integration of a linearized plasmid construct carrying a unique combination of two molecular barcodes. Upon recircularization one barcode remains integrated into the host chromosome, while the other remains on the plasmid allowing direct observation of expansion and contraction of adaptive lineages. We also performed a pilot evolution experiment that allowed us to assess the barcode complexity and establish an analytical framework for the analysis of siBar data.

## Introduction

Evolution of multicopy plasmids occurs at two distinct levels—the intracellular and that of a population (Rodríguez-Beltrán et al. 2021). Mutations occurring during plasmid replication render a bacterial cell carrying such a mutated copy heteroplasmic: immediately after the mutation event the affected cell contains one mutated plasmid and *n*-1 wildtype plasmids. Subsequent rounds of plasmid replication will change this ratio and the fate of the mutation will ultimately be governed by the copy-number control mechanism and its effect on the replication probability for a given plasmid (Paulsson 2002). This is the intracellular level of plasmid evolution in which a mutated plasmid competes against wildtype molecules for the ability to replicate. The effect of the mutation, such as, for example, altering the expression of an antibiotic resistance gene or the potency of its product, will also have an impact on the fitness of the host cell. This is the population level of plasmid evolution where the fate of plasmid mutations ultimately depends on their effect on the host’s fitness. Continuous division of bacterial cells makes the interplay between these two levels of evolution complex. In particular, high copy plasmids without an active partitioning mechanism are distributed randomly among the daughter cells upon division. Due to this “segregational drift” (Ilhan et al. 2019), the ratio of mutated-to-wildtype plasmids is changing constantly and unpredictably (Bedhomme et al. 2017). The segregational drift appears to be a defining factor in restricting the rate of genetic innovation in plasmids lowering the probability of fixation for adaptive mutations (Garoña et al. 2021). What does it take for a mutation to overcome the barrier created by segregational drift and become fixed?

To answer this question it is necessary to identify emerging mutations and trace them through time. Previously, we have applied duplex sequencing to identify emerging adaptive changes in a plasmid conferring antibiotic resistance to *Escherichia coli* cells maintained in a continuous turbidostat culture (Mei et al. 2019). This work showed the emergence of nucleotide changes affecting the copy number control mechanism. The changes likely increased antibiotic resistance of the host by increasing plasmid copy number (and thus increasing the overall expression of the antibiotic resistance gene). However, because we used population sequencing—an approach that does not distinguish individual lineages—we could not visualize the full picture of the underlying dynamics and trace individual mutations. In particular, while the changes were not in genetic linkage we did not know if they appeared sequentially within a single lineage or a small subset of lineages. It was also unclear if the changes appeared independently in several lineages. Finally, we did not know if the changes were maintained in heteroplasmic or homoplasmic state within each adaptive lineage. Answering these questions would require combining sequencing with a lineage tracing approach allowing us to know the identity of evolving clones.

To address these issues we have developed siBar—an experimental framework to **<u>si</u>**multaneously **<u>bar</u>**code chromosomes and plasmids at the lineage-specific level. In any host cell, the single copy of chromosome carries one random DNA barcode, and all intracellular plasmid copies carry another, and identical, DNA barcode. The chromosome barcode is unique among individual cell lineages, so is the plasmid barcode, thus the chromosome/plasmid barcode association is also unique. In this report we describe a pilot experiment showing feasibility of the method, assessing the complexity of resulting barcode libraries, and discussing potential applications.

## Results

### Overview of the method

The siBar rationale is to establish permanent chromosome/plasmid association by introducing unique pairs of chromosomal/plasmid barcodes. This is done by integrating a construct containing a linearized plasmid and two barcodes into the chromosome, and further excising the plasmid and one barcode in such a way that the other barcode remains on the chromosome (Fig. 1). The integration step is referred to as “plasmid-in”, which is mediated through homologous recombination at upstream and downstream homologies using the λ Red system (Datsenko and Wanner 2000). The excision step is referred to as “plasmid-out”, which occurs via recombination at the two FRT (FLP recognition loci) sites flanking the integrated construct (Datsenko and Wanner 2000). The recombination between the two FRT sites is mediated by the site-specific recombinase FLP (Turan and Bode 2011). “Plasmid-out” is a plasmid recircularization process, and the recircularized plasmid is self-replicable. As shown in Fig. 1, two barcodes are designed as flanking the FRT site on the left of the integrated construct. After “plasmid-out”, one barcode is excised together with the plasmid and serves as the plasmid barcode. The other one remains on the chromosome and serves as the chromosomal barcode.

**Figure 1.**
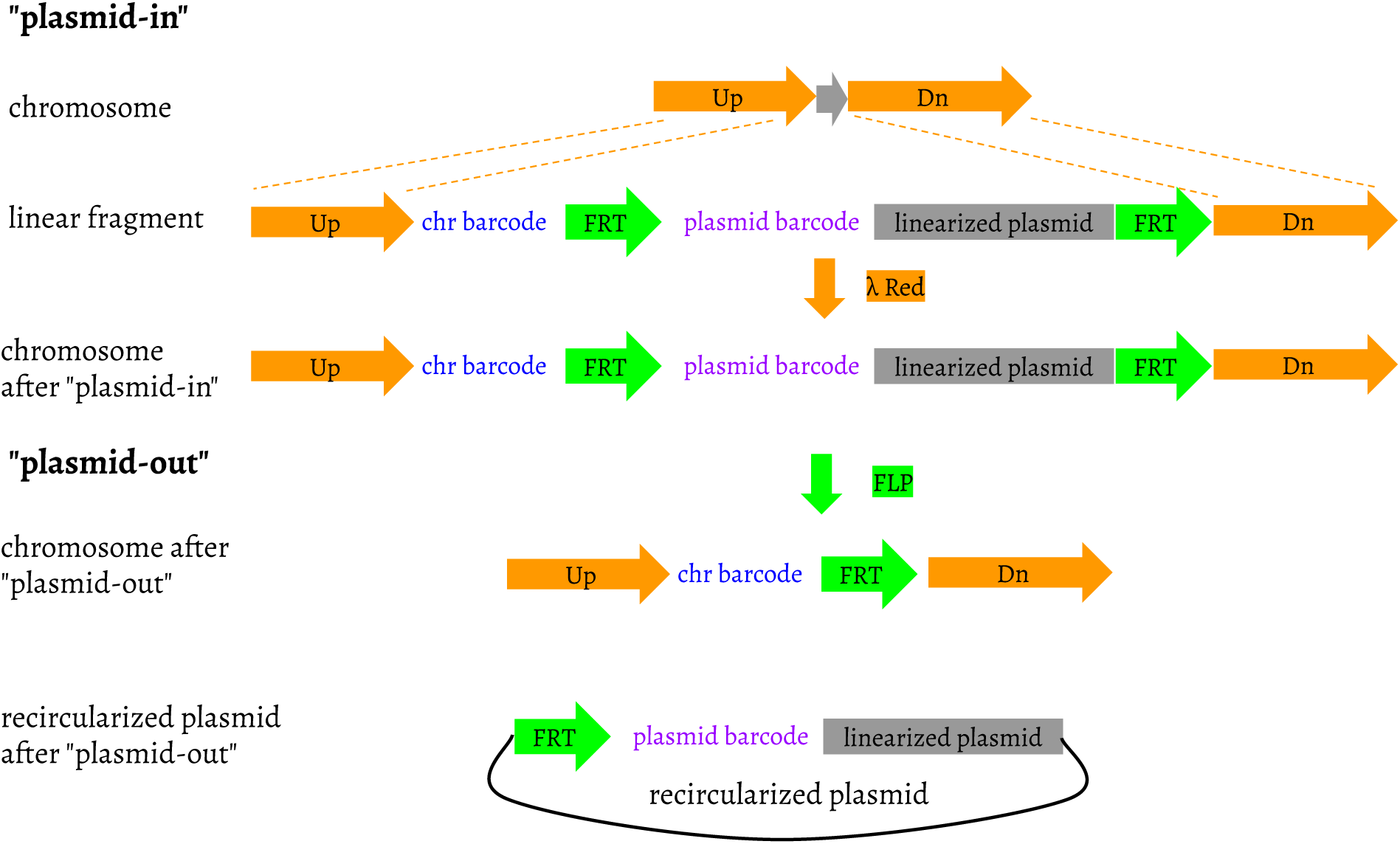

### *galK* locus for plasmid integration and excision

siBar would require a selection/counter-selection strategy to screen recombinants successfully undergoing the “plasmid-in” process, as well as recombinants after “plasmid-out”. Many selection/counter-selection chromosomal loci and cassettes have been reported, including *galK* (Warming et al. 2005), *thyA* (Wong et al. 2005), *tolC* (DeVito 2008), *rpsL-neo* (Zhang et al. 2003), *tetA-sacB* (Li et al. 2013), *cat-sacB* (Lee et al. 2001), and the toxin-antitoxin system (e.g., *ccdA-ccdB* (Wang et al. 2014)). We chose *galK* which catalyzes the first step of the galactose utilization pathway (Csiszovszki et al. 2011). A benefit of using *galK* is that it allows easy screening on Gal indicator plates: *galK* strains show the red Gal+ phenotype, while Δ*galK* strains show the white Gal-phenotype. A conventional use of *galK* as the selection/counter-selection marker involves the galactose analog 2-deoxy-galactose (Warming et al. 2005). We modified the use of *galK* as follows: for a strain (*E. coli* p3478 in this work) carrying an intact *galK*, as shown in Fig. 1, *galK* was first disrupted during “plasmid-in”, leading to the Gal-phenotype; after “plasmid-out”, a chimeric chromosomal *galK* structure was formed, which restored the Gal+ phenotype.

We constructed a series of *galK* fusion plasmids (Fig. 2, plasmid maps in supplementary material) to mimic the chimeric *galK* structure after “plasmid-out”, and tested their Gal phenotypes by transforming them into the Δ*galK* strain JW0740 of the KEIO collection (Baba et al. 2006). Each of these plasmid constructs had a *galK* upstream homology covering a few of its N-terminal amino acid sequences, “C-FRT-G”, and a *galK* downstream homology covering the remaining amino acid sequences till its stop codon. FRT is 34 bp in length. We put a “C” ahead of FRT and a “G” after it, and the “C-FRT-G” structure ensured the in-frame translation of *galK*. We found that at least 6 amino acids before “C-FRT-G” and His21 till the stop codon were necessary for the Gal+ phenotype. His21 and subsequent amino acids have been observed comprising the β1 sheet in GalK structure and the β1 sheet was commonly found in prokaryotic organisms, indicating that β1 could be essential for GalK functionality (Hartley et al. 2004). In contrast, the N-terminal amino acid sequence before His21 varied in length and composition (Hartley et al. 2004). We decided that the chimeric *galK* structure after “plasmid-out” should contain the sequence encoding the first 9 amino acids, and the sequences encoding His21 till its stop codon. To implement this construct, “plasmid-in” needs to be carried out between Ser9 and His21 in *galK*, that is to say, to replace the DNA sequence encoding from Leu10 to Thr20. We have evidence proving that “plasmid-in” at these locations disrupted *galK* and displayed the Gal-phenotype, which is described in *A step-wise plasmid integration procedure*. Note that the chimeric *galK* structure after “plasmid-out” contained more auxiliary elements beyond “C-FRT-G”, which is also shown in *A step-wise plasmid integration procedure*.

**Figure 2.**
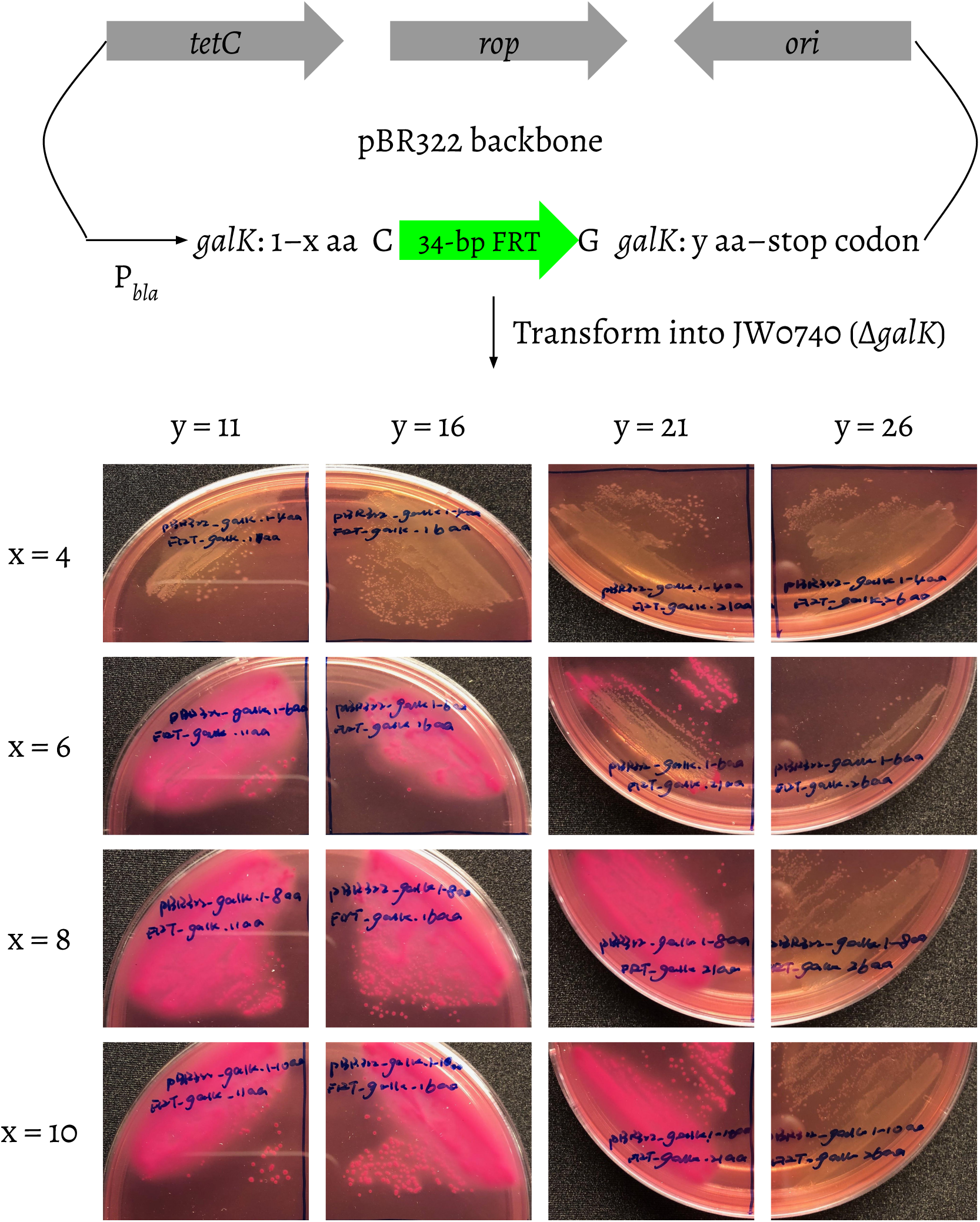

### Conditional control of *polA* expression

The fact that we were trying to integrate the pBR322/ColE1 origin into the chromosome of a commonly used *E. coli* strain would have a lethal effect on the cell due to excessive replication initiation events (Saarilahti and Tapio Palva 1985). For siBar development, we chose *E. coli* p3478 (de Lucia and Cairns 1969) as the host strain. As a derivative of *E. coli* W3110 *thy-*, p3478 bears an amber mutation in *polA*, termed *polA1* (Gross and Gross 1969). *polA1* has the codon of Trp342 mutated from TGG to TAG, leading to a pre-terminated translation of DNA polymerase I (Joyce et al. 1982). This *polA1* mutant has been observed as not being able to support ColE1 replication (Kingsbury and Helinski 1970), and it would allow the pBR322 origin to be integrated into the chromosome. Our solution was to incorporate an additional copy of *polA*, which remained inactive when integrated together with the linearized pBR322 into p3478, but turned active to support pBR322 replication after the plasmid excision (Fig. 3). To make sure the integrated copy of *polA* remained inactive on the chromosome, we constructed three plasmids (Fig. 4, plasmid maps in supplementary material) mimicking the *polA* and *galK* chromosomal structure after “plasmid-in” (Fig. 3). On these plasmids, *polA* and *galK* were put in opposite directions, and were separated by auxiliary sequences. Details about the auxiliary sequences could be found in *A step-wise plasmid integration procedure* (see Methods). By transforming these three plasmids into p3478, we found that if the start codon of *polA* and a ribosome bind site (RBS) were kept ahead of *polA*, *polA* would be active and support the pBR322 origin to replicate in the p3478 *polA1* background, indicating potential transcriptional and/or translational elements in the reverse complementary sequences of *galK*. *polA* would be inactive if the start codon and RBS were removed. As shown in Fig. 3, we decided to integrate the start codon excluded *polA* on the *galK*: 21 aa–stop codon side, and its start codon—ATG, RBS, and the promoter sequences on the *galK*: 1–9 aa side. After “plasmid-out”, we have observed that the plasmid recircularization process turned *polA* active by bringing *polA*, the promoter, the start codon and RBS together. The ability of the recircularized plasmid to self-replicate was confirmed and described in *Gal phenotype confirmation and recircularized plasmid enrichment*. Note that the promoter region of *bla* on pBR322 consists of two promoters in tandem—P1 and P3 (Stüber and Bujard 1981). We only used P1 to avoid *polA* overexpression.

**Figure 3.**
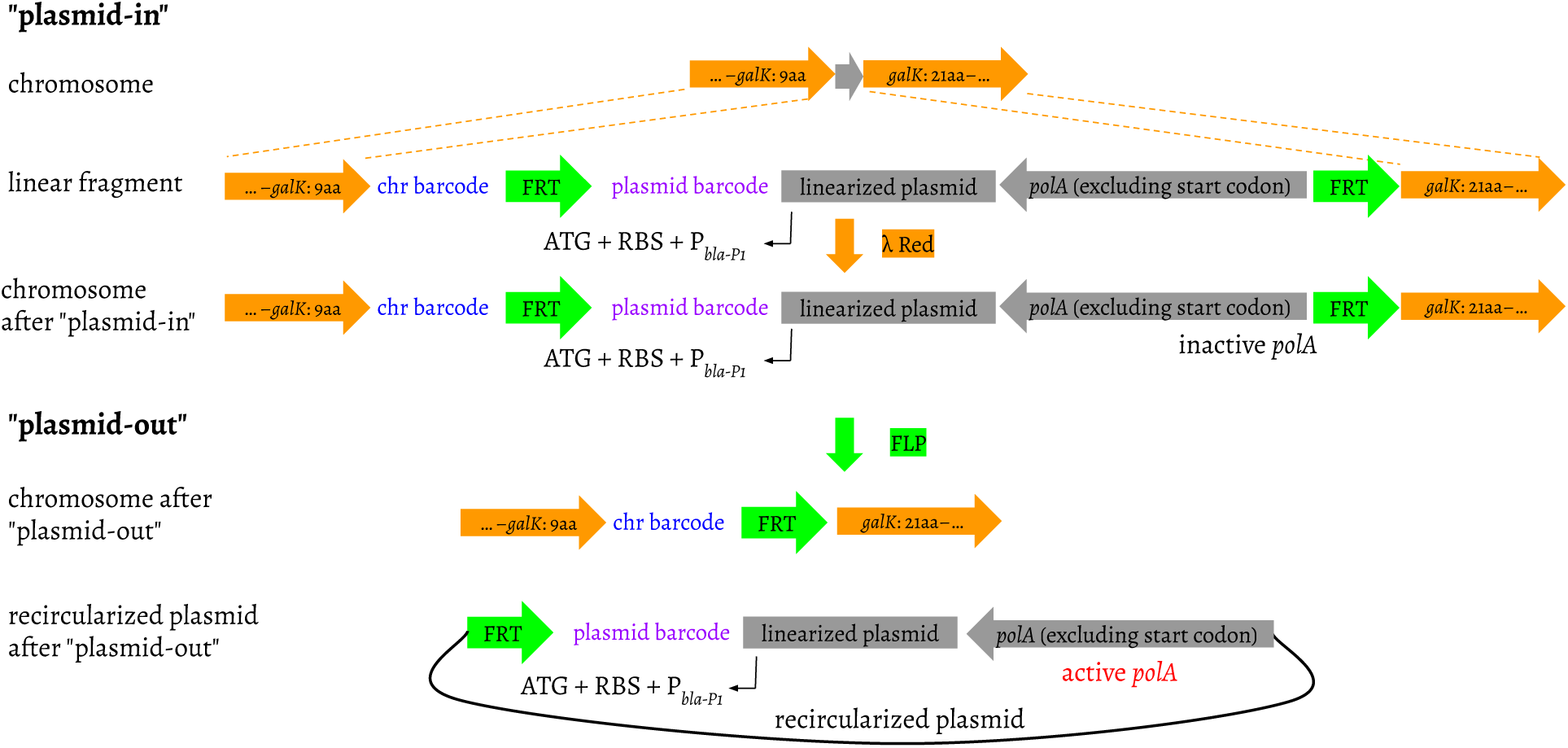

**Figure 4.**
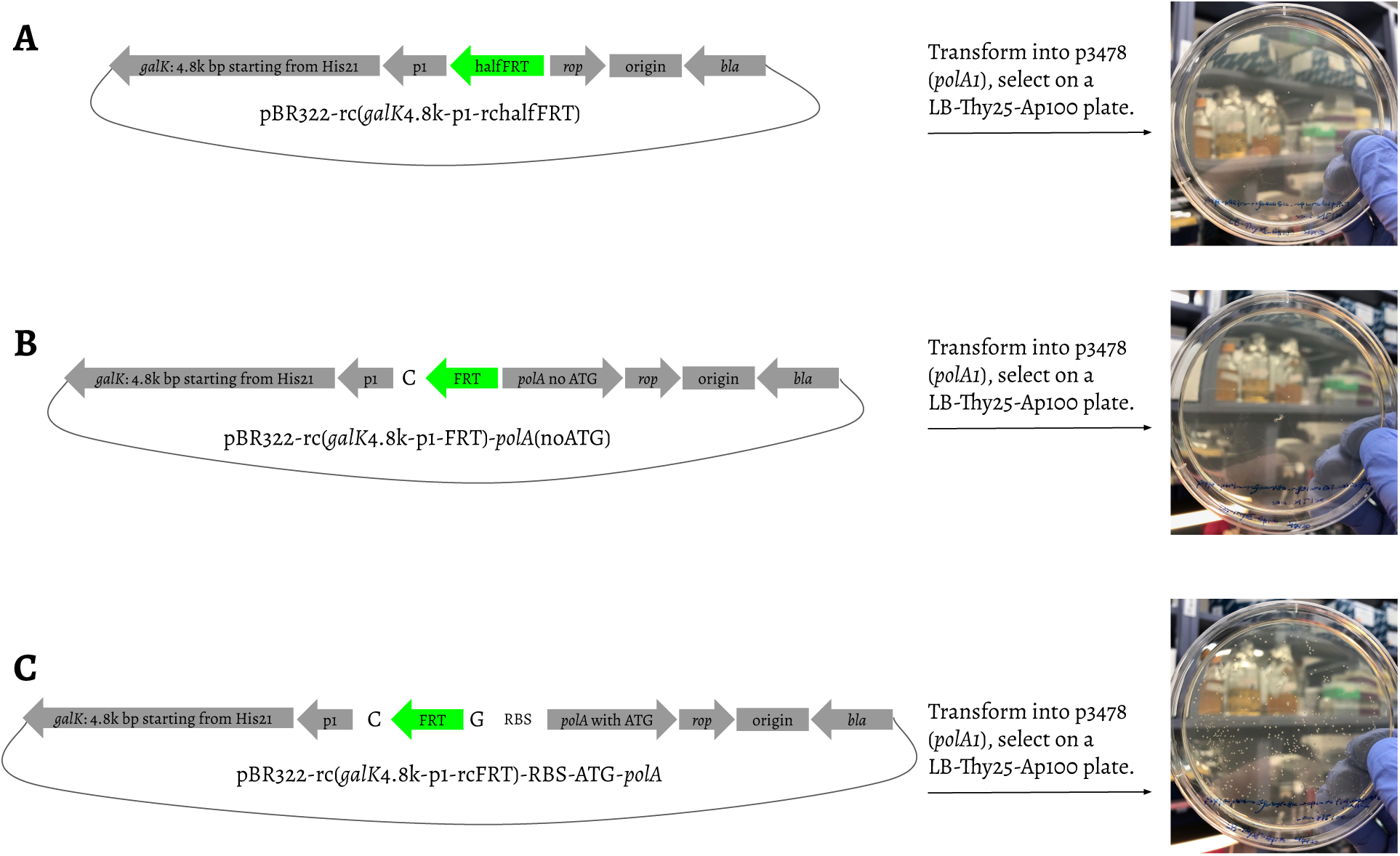

### A step-wise plasmid integration procedure

“Plasmid-in” was divided into five sequential insertion steps, because shorter inserts tend to exhibit higher recombination efficiency (Kuhlman and Cox 2010). pSIJ8 was used as the helper plasmid expressing λ Red for homology based recombination and FLP for FRT based recombination (Jensen et al. 2015; Jensen and Nielsen 2018). *polA* as well as the pBR322 parts—origin, *rop*, and *tetC*—were integrated into the p3478 chromosome in five steps inserting noATG-*polA*5’, *polA*3’, *ori, rop*-*tetC*3’, and barcode-*tetC*5’, respectively. These steps are described in detail in the Methods section. Note that in “plasmid-in” step 5, we isolated three colonies—col1, col2, and col3, and the corresponding cell cultures were referred to as FLP uninduced cultures. Plasmid excision was carried out in these three cultures as described below in the Recircularization section. “Plasmid-in” step 5 and “plasmid-out” step 6 (see below in the Recircularization section) were schematically shown together in Fig. 5 for easy comparison of the chromosomal and plasmid structures before and after the plasmid recircularization process.

**Figure 5.**
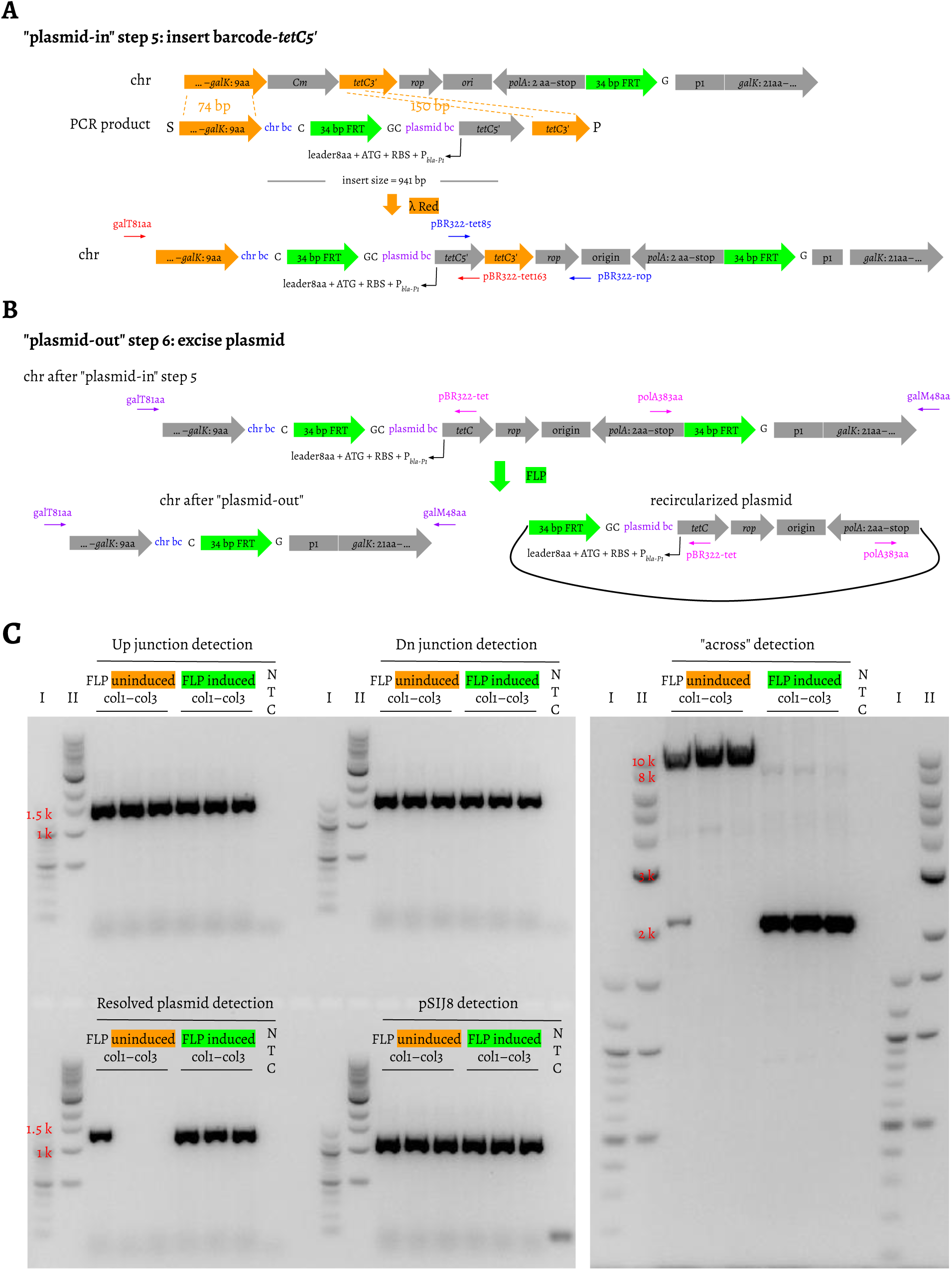

### Recircularization

This step is referred to as “plasmid-out” step 6: excise plasmid. After this step, the whole region flanked by the two FRT sites was excised from the chromosome and recircularized as a single-copy extrachromosomal plasmid carrying the pBR322 origin (Fig. 5B). The auxiliary nucleotides introduced in “plasmid-in” step 5 ensured the in-frame translation of *galK* and *polA* after “plasmid-out” step 6 (Fig. 5B). The chromosomal and plasmid barcodes were designed as 21 bp random nucleotides—7 × “VNN”. “V” stands for “A/C/G”, which avoids the appearances of stop codons in “VNN”, since the chromosomal and plasmid barcodes were translated with *galK* and *polA* respectively. After plasmid excision, the chromosomal *galK* frame would be: *galK*: 1–9 aa, chromosomal barcode (21 bp), “C-FRT-G” (36 bp), p1 (18 bp), and *galK*: 21 aa–stop codon. The plasmid *polA* frame would be: “ATG”, a leader peptide sequence (24 bp), plasmid barcode (21 bp), the reverse complementary sequence of “FRT-GC” (36 bp), and *polA*: 2 aa–stop codon.

During “plasmid-out” step 6, plasmid excision was performed by inducing FLP expression in the FLP uninduced cultures of the three isolated colonies in “plasmid-in” step 5, which led to the FLP induced cultures. We did not perform colony purification on the FLP induced cultures, thus the FLP induced cultures were mixtures of cells having experienced the FLP/FRT recombination or not. As shown in the lanes marked under “FLP induced” in Fig. 5C, the PCR products using the primer pair galT81aa and galM48aa (Fig. 5B) were 2,136 bp. The shift from 8,465 bp to 2,136 bp comparing the FLP uninduced to the FLP induced indicates the plasmid excision. We also observed a faint 2,136 bp band for the FLP uninduced col1, indicating a FLP expression leakage from the helper plasmid pSIJ8 (Jensen et al. 2015). We used another primer pair—pBR322-tet and polA383aa (Fig. 5B)—targeting the recircularized plasmid to further confirm plasmid excision. No PCR products would be seen before plasmid excision, and 1,400 bp PCR products could be detected after plasmid recircularization. In the FLP induced, the 1,400 bp bands were detected in all of three colonies. This band was also seen in the FLP uninduced col1, which is consistent with the observation of the pSIJ8 FLP expression leakage. The 2,136 bp products generated by galT81aa and galM48aa and the 1,400 bp products generated by pBR322-tet and polA383aa amplifying the FLP induced culture of col1 were subjected to Sanger sequencing (Fig. 6, B and C). The barcode sequence compositions were observed identical prior to (Fig. 6A) and after plasmid excision (Fig. 6, B and C). The Sanger results also confirmed that the plasmid barcode was excised together with the plasmid, and the chromosomal barcode remained on the chromosome.

**Figure 6.**
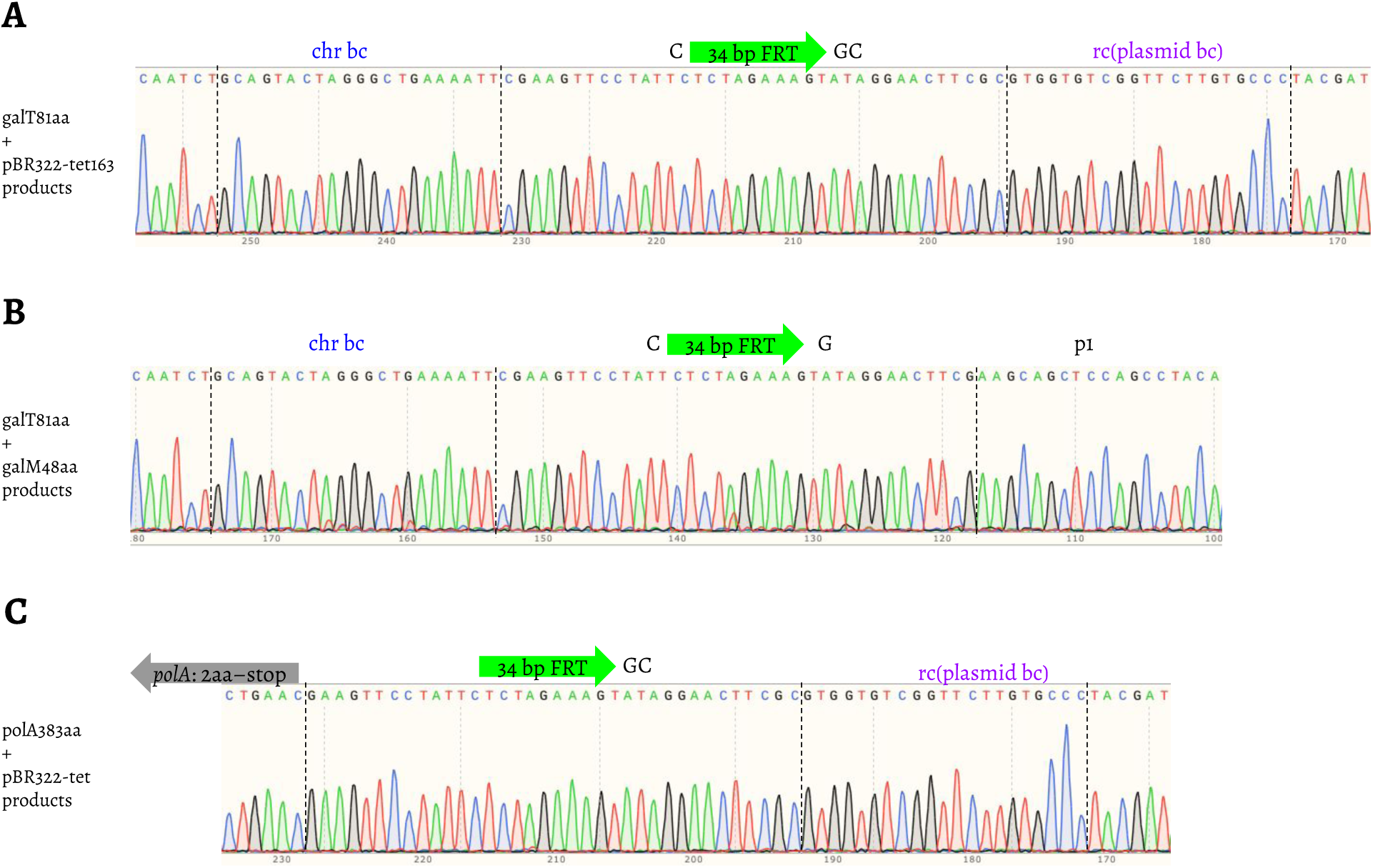

### Gal phenotype confirmation and recircularized plasmid enrichment

The FLP uninduced and induced cultures of the three colonies from “plasmid-in” step 5 and “plasmid-out” step 6 were streaked on the minimum M63 agar plates with galactose as the sole carbon source (Fig. 7A, first column top plate). The FLP uninduced cannot grow, but the FLP induced can. This suggests that *galK* was disrupted after “plasmid-in” step 5 and showed the Gal-phenotype. The chimeric *galK* structure after “plasmid-out” step 6 restored the Gal+ phenotype. We also streaked the parent strain p3478, p3478 transformed with pSIJ8, and the resulting strains of “plasmid-in” steps 1–4 on the same M63 galactose agar plates (Fig. 7A, first column bottom plate). p3478 and p3478 with pSIJ8 can grow, but the “plasmid-in” steps 1–4 resulting strains cannot, which supports our earlier claim that *galK* has been disrupted in the “plasmid-in” steps 1–4.

**Figure 7.**
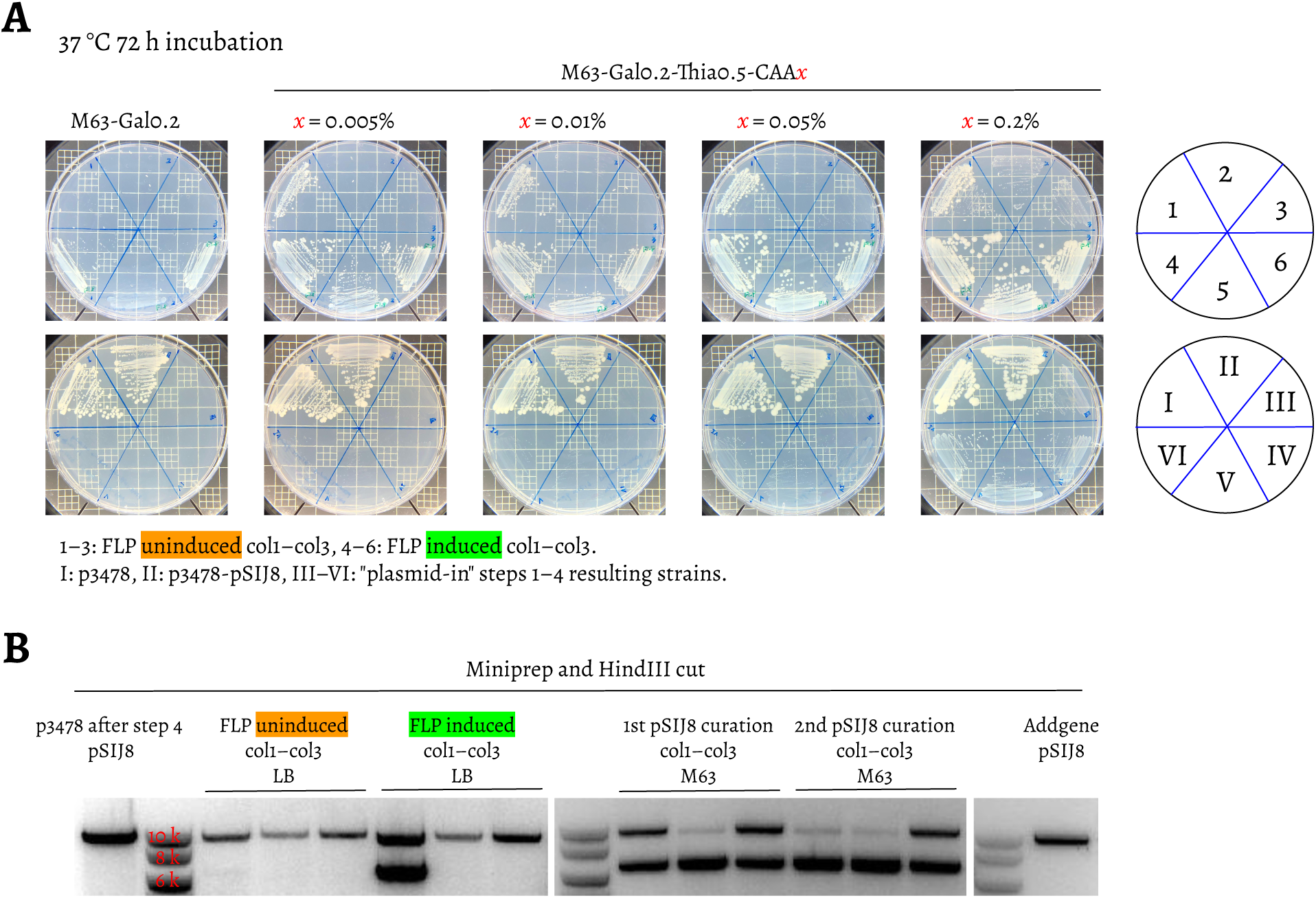

We have observed slow and different growth rates for the strains mentioned above on M63 plates with galactose. Thiamine and casamino acids (CAA) were then supplemented to support their growth. Since CAA has been reported as being able to serve as a carbon source (Farrell and Finkel 2003), we tried to supplement CAA at different concentration levels—0.005%, 0.01%, 0.05%, and 0.2%. As shown in Fig. 7A (second to last columns), at CAA concentrations 0.05% and lower, the growths of the FLP induced were improved compared to those without thiamine and CAA supplemented. For the FLP uninduced, only col1 was able to grow, which is consistent with the FLP leakage expression observed in col1. At CAA concentration 0.2%, the growths of the FLP uninduced of col2 and col3 became fairly visible. As we increased the CAA concentration, the resulting strains of “plasmid-in” steps 1–4 also started to grow. These results suggest that CAA can serve as a carbon source indeed, and needs to be used at concentrations 0.05% or lower.

After plasmid excision, each of the three colonies contained two types of extrachromosomal plasmids—pSIJ8 and the recircularized plasmid. To cure pSIJ8 and enrich the recircularized plasmid, we propagated the FLP induced cultures at 37 °C in M63 broth supplemented with galactose, thiamine, 0.05% CAA, and 10 μg/mL Tc. The tolerance of Tc has been reported to linearly increase with the *tetC* copy number (Gutterson and Koshland 1983). Before *tetC* was excised from the chromosome, it was in a single copy form in each cell, and we used Tc at 5 μg/mL. After plasmid excision, we increased the Tc concentration to 10 μg/mL. After two serial passages in the M63 broth, the recircularized plasmids were enriched, and pSIJ8 was gradually cured (Fig. 7B).

### Initial assessment of barcode complexity

We carried out a pilot run by combining ten electroporation reactions as described in “plasmid-in” step 5. As shown in Fig. 8A, the ten reactions were spread on the Gal indicator plates, then the plate surfaces were washed by LB broth and pooled as the FLP uninduced culture. The FLP expression was then induced, which led to the FLP induced culture. The FLP induced culture was passaged in two parallel lines—M63 and LB. In the M63 line, galactose was used as the sole carbon source. Thiamine, 0.05% CAA, and Tc were supplemented. We first grew the FLP induced into M63 with 10 μg/mL Tc, and then 30 μg/mL Tc. In the LB line, the FLP induced was first passaged into 10 μg/mL Tc, followed by increasing Tc concentrations in the order of 10 → 30 → 50 → 100 μg/mL.

**Figure 8.**
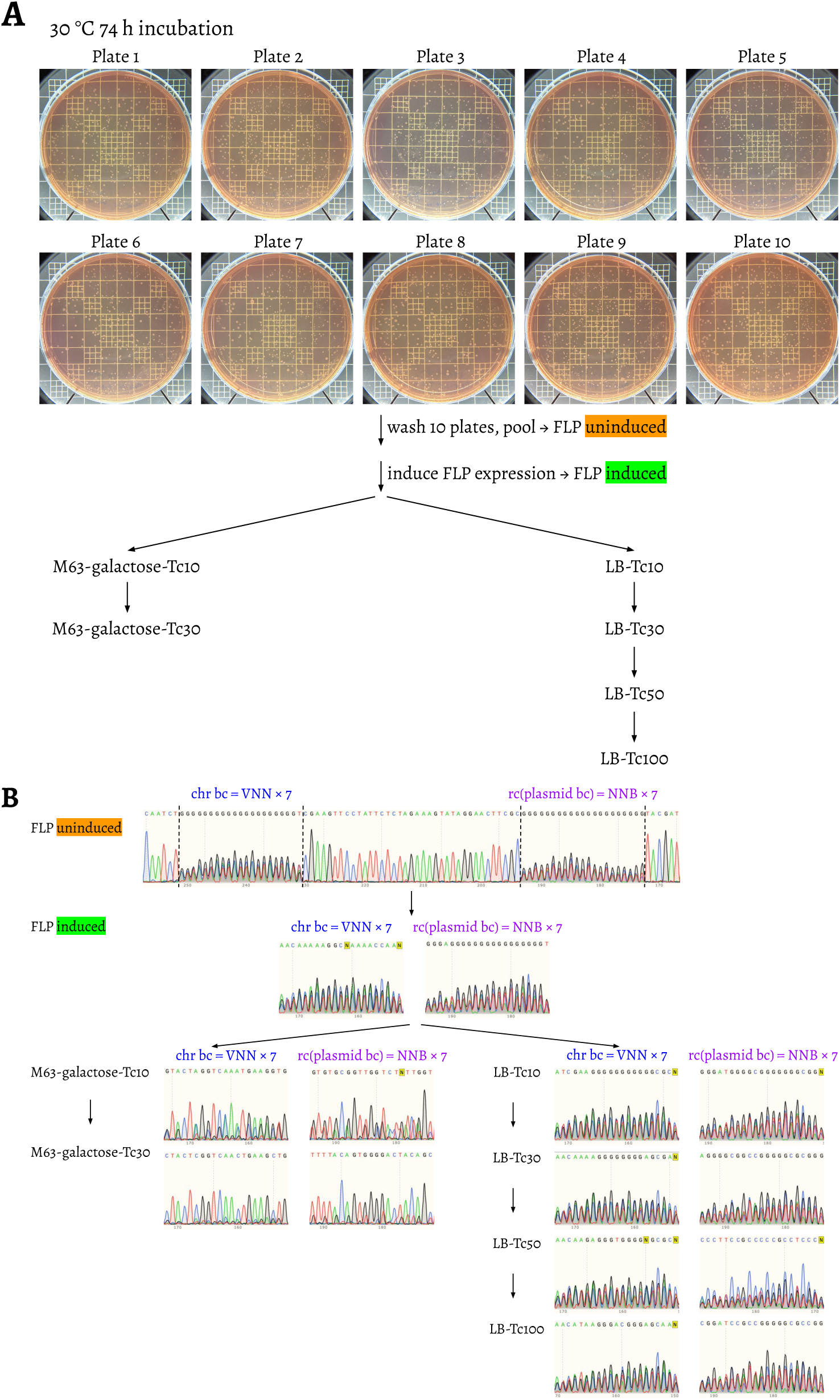

We then performed Sanger sequencing on the barcode regions of the FLP uninduced, FLP induced, and the two parallel lines (Fig. 8B). The barcode diversity was high in the FLP uninduced and FLP induced as indicated by mixed peaks. After transferring into the M63 line, the diversity collapsed in 10 μg/mL Tc, and further decreased in 30 μg/mL Tc. On the contrary, in the LB line, the barcode diversity was well maintained, even when 100 μg/mL Tc was applied.

We chose the following four cell culture samples to perform next-generation sequencing: (1) the FLP uninduced sample, (2) the FLP induced sample, (3) the M63 line sample under 10 μg/mL Tc, and (4) the LB line sample under 100 μg/mL Tc (Fig. 9). The ampliconic libraries were prepared using a two-step PCR protocol modified from that described by Levy et al. (Levy et al. 2015). These amplicons were generated by first performing a 3-cycle PCR to amplify the barcode region using forward and reverse primers containing unique molecular identifiers (UMIs). One template DNA molecule would only generate one product molecule after the 3-cycle PCR. The UMIs will be used to remove biases in molecule counts—referred to as PCR jack-potting—in downstream analyses. The 3-cycle PCR products were purified and used as the templates for the second round of multiple-cycle PCR using the Nextera index primers. For the FLP uninduced sample, both the chromosomal and the plasmid barcodes were on the chromosome, thus there would be only one amplicon covering the two barcode regions (Fig. 9A). For the remaining three samples, each would generate two replicons—one for the chromosomal barcode region (Fig. 9B), and another for the plasmid barcode region (Fig. 9C). Seven sequencing library samples were generated in total.

**Figure 9.**
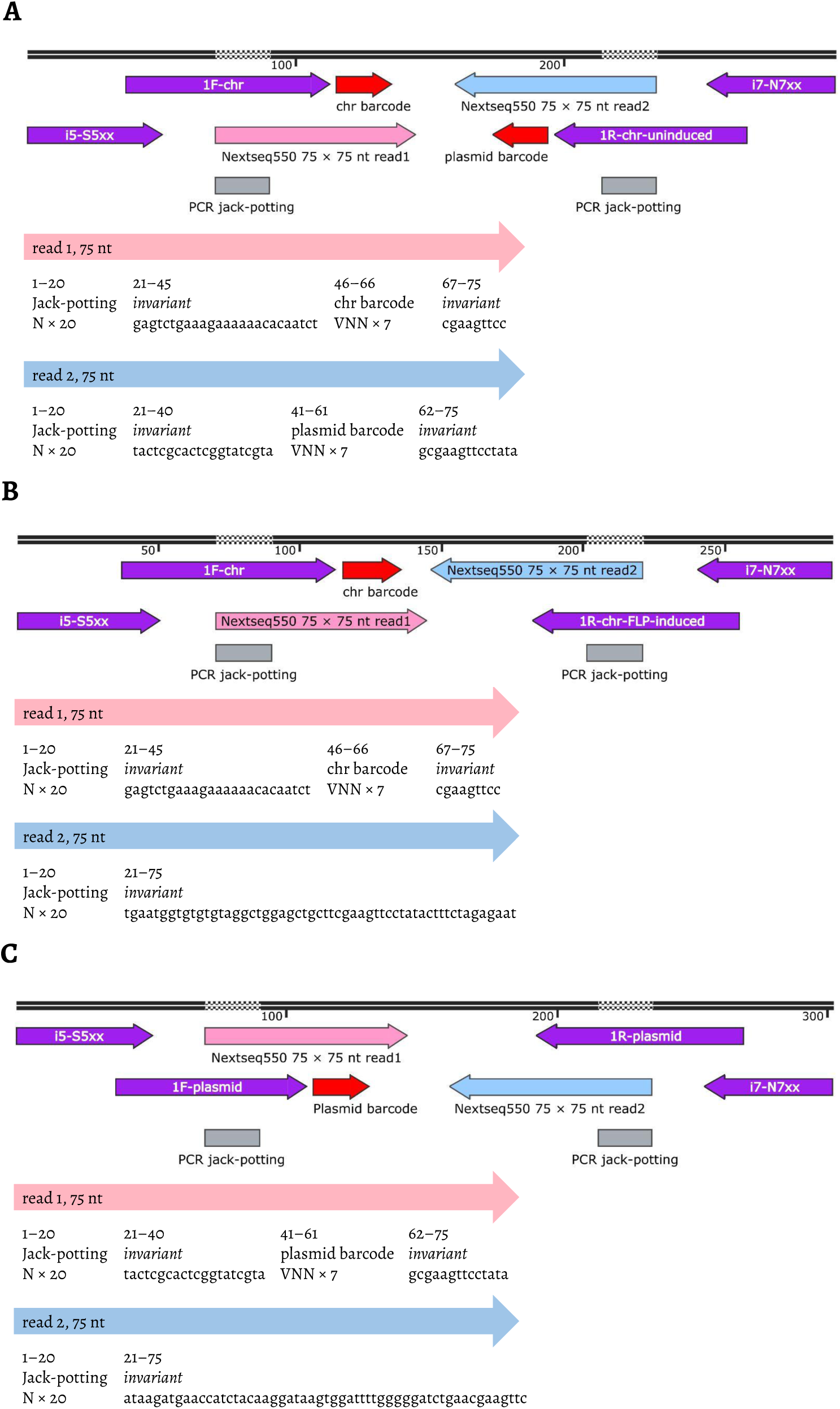

In the FLP uninduced sample, we identified 2,952 lineages with unique chromosomal and plasmid barcodes combinations (Figs. 10 and 11, Table S3). In the FLP induced samples, 1,243 chromosomal and 774 plasmid lineages were recovered respectively from the pool of the 2,952 lineages; 739 lineages were found with both the chromosomal and the plasmid barcodes detected (Figs. 10 and 11). In the M63 line sample, 543 chromosomal and 377 plasmid lineages were found, and 330 lineages were found with both barcodes detected (Fig. 10). In the LB line sample, 798 chromosomal and 719 plasmid lineages were found respectively, and 651 lineages were identified with both barcodes detected (Fig. 11). There were fewer lineages detected in the M63 line sample compared to the LB line sample, which is consistent with the Sanger sequencing results (Fig. 8B) showing a lower barcode diversity in the M63 line sample. We reasoned that the chromosomal barcode (VNNs) after “plasmid-out” might have an effect on galactose utilization since the barcodes were translated in-frame within the chimeric *galK* structure. In LB broth, the carbon sources are mostly catabolizable amino acids, with a low concentration of sugars (Sezonov et al. 2007).

**Figure 10.**
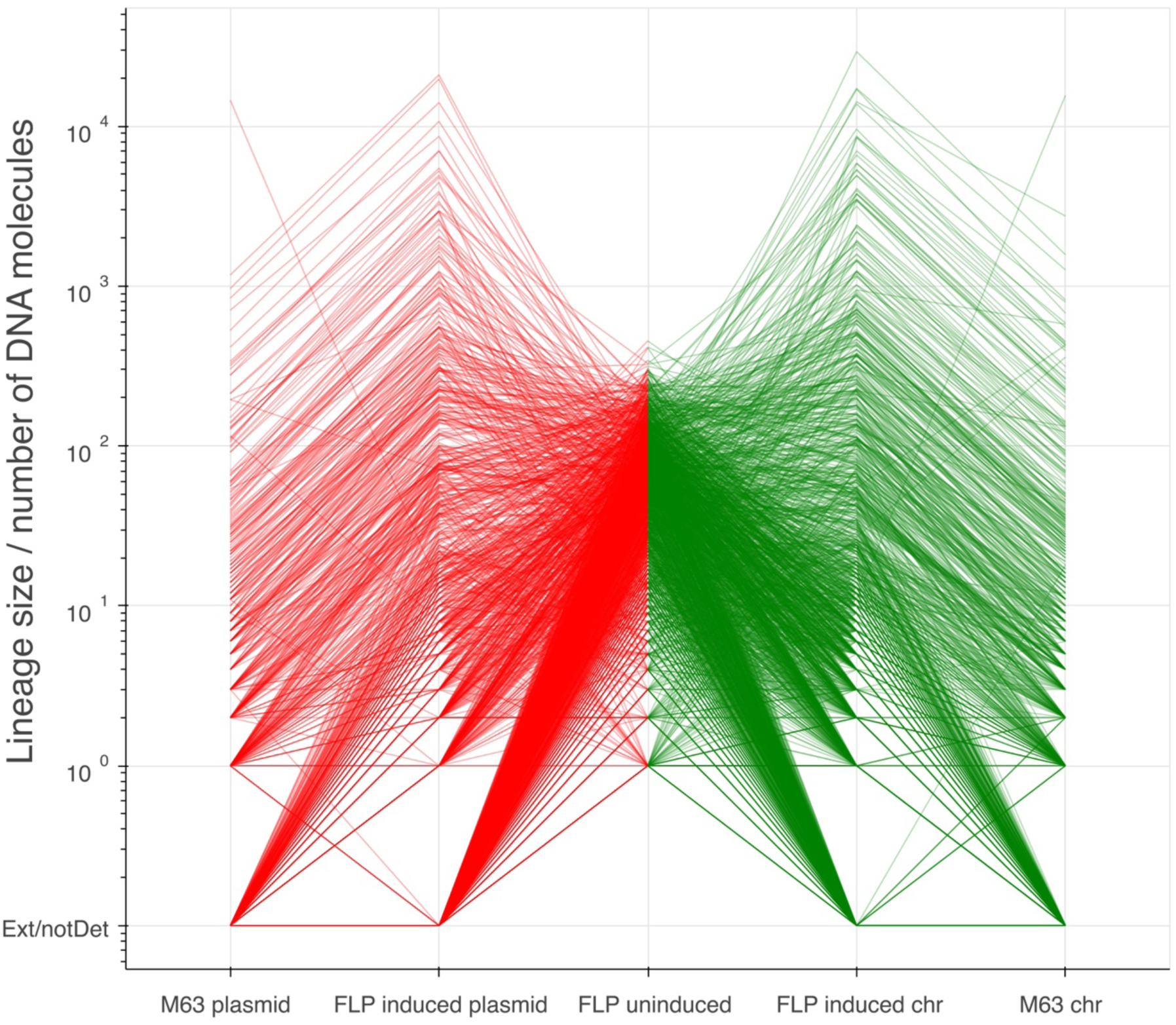

**Figure 11.**
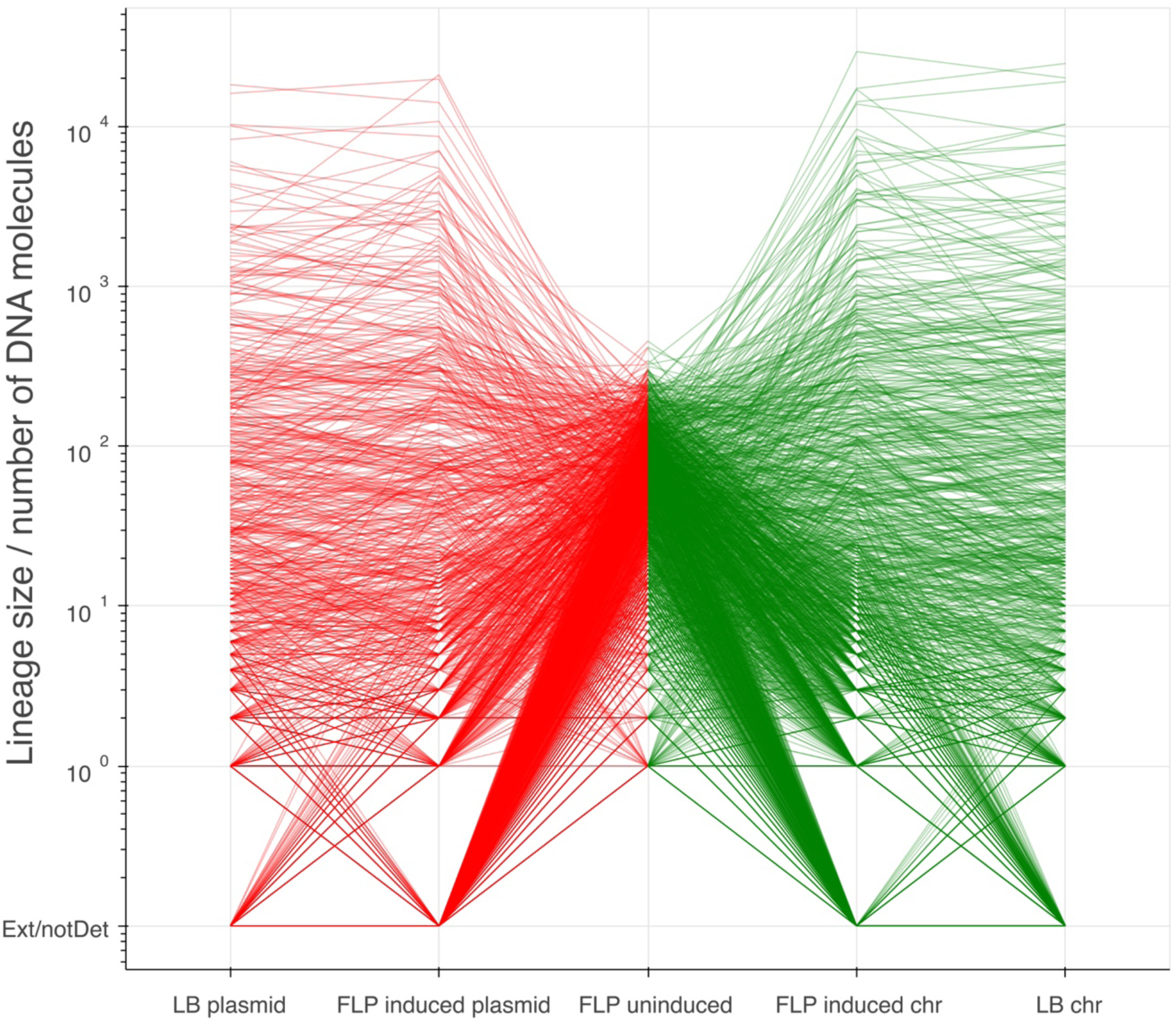

We have observed two types of discrepancies in the number of lineages identified. The first discrepancy is the number of lineages detected in different cell culture samples, e.g., 2,952 vs. 1,243 chromosomal lineages comparing the FLP uninduced and induced samples. This type of discrepancy is mostly due to the differences in lineage size. For simplicity, it should be noted that we are not considering sampling errors here. Also, the number of reads from the Nextseq 550 run allocated to each of the seven ampliconic libraries was not exactly the same. We define the lineage size as the number of DNA molecules in a lineage. For example, in the FLP uninduced library, the largest lineage has 455 chromosomal DNA molecules, and in the FLP induced chromosome library, the largest lineage has 29,341 chromosomal DNA molecules (Table S3). For each of these two libraries, approximately the same amount of DNA was used as the two-step PCR template (Fig. 12A). The amount of template DNA was the product of the number of lineages and the lineage sizes. Given that the lineage sizes in a library are larger (e.g., comparing lineage size: FLP induced chromosome > FLP uninduced), fewer lineages would be included in the library (e.g., comparing number of lineages: FLP induced chromosome < FLP uninduced). The second discrepancy is the number of chromosomal and plasmid lineages detected in the same cell culture sample, e.g., 1,243 chromosomal and 774 plasmid lineages were detected in the FLP induced chromosome and plasmid libraries. This discrepancy indicates the plasmid copy number heterogeneity (Wong Ng et al. 2010; Jahn et al. 2016). If the plasmid copy number were identical in individual cells, let us assume a plasmid copy number of 2 as shown in Fig. 12B, the amount of plasmid molecules would be twice as many as that of chromosome molecules. Assuming the same amount of chromosome and plasmid molecules were used as the template of each ampliconic library, the numbers of chromosomal and plasmid lineages identified should be equal and unaffected by the universal doubling process as shown in Fig. 12B (again, assuming no sampling error). However, if the plasmid copy numbers varied in different cells, the numbers of chromosomal and plasmid lineages captured would be different (Fig. 12C).

**Figure 12.**
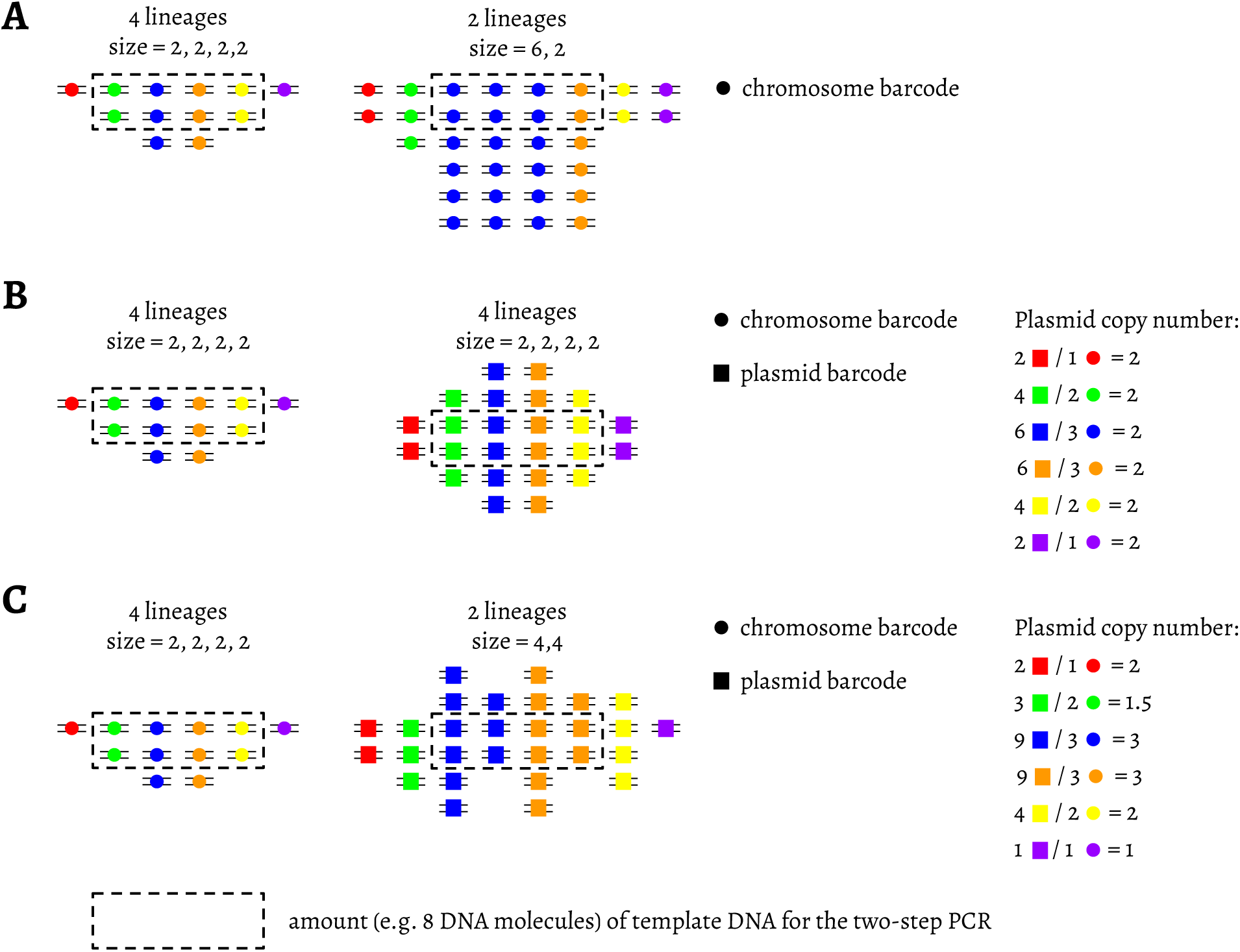

## Discussion

In this paper we describe a method for simultaneous tracking of genome/plasmid associations. One potential limitation of our approach is that it does not allow direct observation of plasmid dynamics at the single cell level. In other words, if we detect a mutation associated with a plasmid barcode while we also observe wild type plasmids with the same barcode, we do not know whether they are contained within the same cell or not. To overcome this limitation we will attempt to use plasmid copy number estimation as a proxy for identifying heteroplasmic states. Fig. 13 illustrates this idea. It shows a portion of a genealogy for a particular chromosomal/plasmid combination starting with a single transformant. While we cannot directly observe whether mutated plasmids (red in Fig. 13) are present in homoplasmic or heteroplasmic states, we can try to infer this from copy numbers (blue boxes). The difficulty with the approach is that once population expands copy numbers for both mutated and wild type plasmids will become high and it will be impossible to infer heteroplasmy given a single sampling point. However, previous results (Bedhomme et al. 2017) point to the fact that the time frame in which mutated and wildtype plasmids co-exist in population are brief. Thus if we detect mixtures of mutated and wild type plasmids sharing a single barcode for a prolonged period of time, we may assume that they are present in heteroplasmic state (Rodriguez-Beltran et al. 2018).

**Figure 13.**
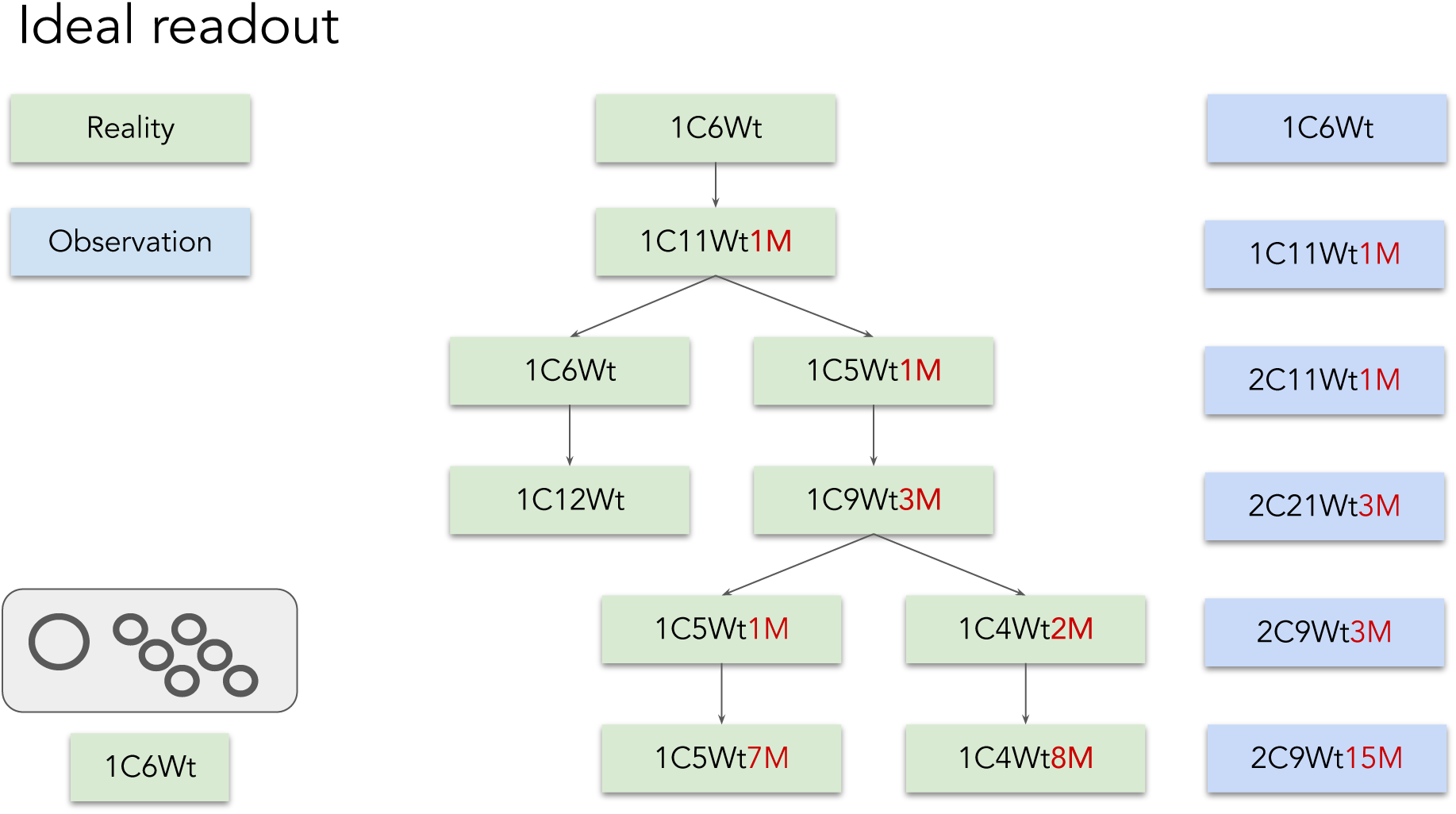

The ability to compute relative copy numbers for genomes and plasmids will require a minor modification to siBar protocol that we are currently implementing. In the original protocol we applied the two-step PCR process to prepare two separate ampliconic libraries—the chromosomal amplicon and the plasmid amplicon. Approximately the same amount of chromosome and plasmid DNA molecules were used as the PCR templates, thus making it inappropriate for plasmid copy number estimation in this current form. To remove this limitation we plan to amplify the chromosome and the plasmid in one PCR reaction, with both of the two sets of primer pairs added, leading to the products of the two amplicons in one reaction. For an individual cell lineage, i.e., a cell lineage carrying its unique chromosomal and plasmid barcodes, by merging the UMIs tagged to individual DNA molecules, we will know the numbers of chromosomes and plasmids present in the template genomic DNA pool before PCR amplification.

Our method can be used to design experimental evolution experiments mimicking “evolutionary games” observed in cases when encounters of animals with antagonistic features within a population do not always escalate to fights (Smith 1982). The cooperative behavior suggests that selection acts on the population level, and is good for a species; this seems to contradict the Darwinian viewpoint that selection acts on the individual level, where individuals with differential fitness are always fighting (Broom and Krivan 2018). We are planning to apply siBar to compare the effects of taking these two selection strategies. In our previous work, we have observed two types of variants on the plasmid pBR322, both of which boosted the antibiotics resistance levels of their host cells by increasing the plasmid copy number, thus achieving higher expression levels of the antibiotics resistance gene (Mei et al. 2019). The difference between these two types of variants is that one type (variants at positions 3,027–3,035) only affected the copy number of the mutant plasmids carrying the variants, thus conferring mutant plasmids differential fitness. The other type (variant at position 3,118) increased the copy numbers of all intracellular plasmids—both the wild-type and the mutant plasmids, which reflected a cooperative behavior. The excised plasmid carries the same ColE1 origin of replication as pBR322, and the same Tc resistance gene. By forcing host cells carrying the excised plasmid to grow under Tc pressure, it is likely that the variants increasing the plasmid copy number would be observed again. By comparing the growth behavior of cell lineages carrying the two types of variants, we will be able to tell which of the two selection strategies is preferred.

## Materials and Methods

### Strains, media and chemicals

*E. coli* strain DH5α was obtained from NEB (cat. C2987I). *E. coli* p3478 (W3110 *thy-polA1*, cat. 4303), the KEIO collection Δ*galK* mutant strain JW0740 (cat. 8803), and BW25141 (*pir*+, cat. 7635) were obtained from *E. coli* Genetic Stock Center (CGSC) at Yale University. LB broth was made according to the manufacturer’s instructions (EMD cat. 110285). Gal indicator plates were made using MacConkey Agar Base (BD cat. 281810) supplemented with 0.2% galactose (Sigma-Aldrich cat. G5388). 2× M63 buffer was made by adding 3 g KH_2_PO_4_ (Sigma-Aldrich cat. P5655), 7 g K_2_HPO_4_ (Sigma-Aldrich cat. P8281), 2 g (NH_4_)_2_SO_4_ (Sigma-Aldrich cat. A4418) to 500 mL H_2_O, followed by autoclaving. M63 borth with 0.2% galactose was made by diluting 500 mL 2× M63 buffer to 1× using 500 mL H_2_O and adding filter sterilized 0.1 mL 5 mg/mL FeSO_4_ · 7H_2_O (Sigma-Aldrich cat. F8633), 1 mL 1 M MgSO_4_ (Sigma-Aldrich cat. M2773), and 20 mL 10% galactose. All media was supplemented with 0.05% antifoam B (Sigma-Aldrich cat. A5757). For p3478 and derivative strains, thymine (Sigma-Aldrich cat. T0895) was added to the media at the working concentration of 25 μg/mL. For template plasmid cloning purposes, ampicillin (Ap, Sigma-Aldrich cat. A9518), kanamycin (Km, Sigma-Aldrich cat. K1876), chloramphenicol (Cm, Sigma-Aldrich cat. C0378), tetracycline (Tc, Sigma-Aldrich cat. T3383) were used at 100, 50, 20, 10 μg/mL, respectively. For recombination purposes, arabinose (Sigma-Aldrich cat. A3256) and rhamnose (Sigma-Aldrich cat. 83650) were used at 15 and 50 mM to induce λ Red and FLP expression on pSIJ8, respectively.

### Plasmids and primers

All plasmids used in this study are listed in Table S1. pBR322 was obtained from NEB (cat. N3033S). The helper plasmid pSIJ8 expressing the λ Red and FLP recombinase was obtained from Addgene (cat. 68122). pSIJ8 can replicate at 30 °C, but not at the nonpermissive 37 °C. The conditional replicon *oriRγ* was cloned from pKD4 obtained from CGSC (cat. 7632). pKD13 (CGSC cat. 7633) and pKD3 (CGSC cat. 7631) were used as the templates to clone the Km^R^ and Cm^R^ genes, respectively. Plasmid maps are supplemented in SnapGene Viewer dna format, including the plasmids to explore the *galK* integration site, the plasmids to test potential and/or translational elements in *galK*, and the PCR template plasmids for each step of the “plasmid-in” process. The primers used to construct these plasmids were obtained from IDT and supplemented in Table S2.

### PCR products generation for chromosomal insertion

The PCR products for insertion purposes were generated using the template plasmids in Table S1 and the primers in Table S2. We used two high fidelity PCR systems—Q5 (NEB cat. M0492), and Platinum SuperFi II (Invitrogen cat. 12368010)—for PCR products generation. For the primers used to generate the insertion fragments in “plasmid-in” steps 1–4, each primer contained sequences annealing to the template plasmid, and sequences homologous to the chromosomal insertion sites.

Below we are describing the insertion fragments generation process in “plasmid-in” step 5. We are not showing these processes in “plasmid-in” steps 1–4. They were essentially the same as the first round of PCR in “plasmid-in” step 5, but with different template plasmids (Table S1) and primers (Table S2). Note that they did not require the second round of PCR as in “plasmid-in” step 5.

In “plasmid-in” step 5, the goal was to generate as many colonies/lineages as possible. Two rounds of PCR were performed to increase the homology length and add the 5’-phosphothioated and 5’-phosphorylated modifications to the PCR products. In the first round, the template plasmid was amplified using the oligos *galK9aa-cBc-C-34FRT-GC-pBc-8aa-RBS-EcoRI* as the forward primer and *pBR322-tetCDn150(EagI included)-tetC50* as the reverse primer (Table S2).

The VNNs in the forward primer are the chromosomal barcodes. The NNBs in the forward primers are the plasmid barcodes. The plasmid barcodes will be translated in the reverse complementary frame of the NNBs (B = T/G/C), generating the VNNs plasmid barcodes after plasmid excision.

The first round PCR products were purified by 0.5× beads (Beckman Coulter cat. A63881), and used as the template of the second round of PCR using the oligos *Up_S_ext40_prime_20* as the forward primer and *Dn_P_ext40_prime_20* as the reverse primer (Table S2).

The forward and reverse primers in the second round PCR extended the homology lengths. Moreover, the forward primer was 5’-phosphothioated and the reverse primer was 5’-phosphorylated. The 5’-phosphorothioation modification on the *galK*: 1–9 aa side and the 5’-phosphorylation modification on the *galK*: 21 aa–stop codon side can increase the recombination efficiency (Maresca et al. 2010; Mosberg et al. 2010; Jensen et al. 2015).

The PCR products were 0.5× beads purified, DpnI (NEB cat. R0176) treated, 0.5× beads purified, suspended in H_2_O, quantified by Qubit BR (Invitrogen cat. Q32850), and ready for “plasmid-in” chromosomal insertion.

### Electrocompetent cells preparation and “plasmid-in” recombination

p3478 and derivative strains containing pSIJ8 (Jensen et al. 2015) were grown in LB broth supplemented with 25 μg/mL thymine and 50 μg/mL Ap at 30 °C to the early exponential phase (OD_600_ = 0.2–0.3). Arabinose was then added at a final concentration of 15 mM to induce λ Red expression. The cell culture was allowed extended growth to the mid exponential phase (OD_600_ = 0.4–0.6). Cells were then collected and made electrocompetent by washing twice in ice-cold distilled H_2_O and three times in ice-cold 10% glycerol. ∼2 μg purified PCR products were electroporated into 50 μL electrocompetent cells at 1.75 kV using 0.1 cm cuvettes (Bio-Rad cat. 1652089) and a MicroPulser electroporator (Bio-Rad cat. 1652100). Electroporation reactions were immediately recovered in 1 mL SOC (BD cat. 244310) supplemented with 25 μg/mL thymine, transferred to 14 mL tubes (BD Falcon cat. 352059), and allowed outgrowth at 250 rpm at 30 °C for 2 h. Outgrowth culture was concentrated, spread on LB (“plasmid-in” steps 1–4) or Gal indicator (“plasmid-in” step 5) plates with 25 μg/mL thymine and antibiotics, and incubated at 30 °C. The choice of antibiotics for each of “plasmid-in” steps 1–5 can be found in “plasmid-in” steps 1–5 (Figs. 13–16, and 5). Km, Cm, and Tc were used at 25, 20, 5 μg/mL for screening colonies with integrated chromosomal antibiotic gene markers. Colonies were isolated from the antibiotics agar plates, and purified once on the same plates. After insertions of PCR products, two novel junctions were created at the upstream and downstream recombination homologies. Locus-specific primers (Table S2) flanking each junction were used to verify the recombination at the homology. We also performed PCR amplification using primers covering the following three regions: upstream junction, insert, and downstream junction. PCR products covering the three regions were sequenced by Sanger sequencing to further confirm successful insertions. One colony verified with correct insertions was used as the starting strain for the next “plasmid-in” step. Note that we did not use Ap to help keep pSIJ8 when the outgrowth culture was grown on antibiotics agar plates because of the additional stress that Ap might impose. To confirm the presence of pSIJ8 after Km, Cm, or Tc selection, a region of pSIJ8 was amplified (primers in Table S2).

### Integration procedure

#### “plasmid-in” step 1: insert noATG-*polA*5’

Because *polA* is ∼2.7 kb in length we inserted it in two steps. In “plasmid-in” step 1, ∼1.2 kb of the 5’-end of *polA* (with the start codon excluded) and the kanamycin resistance (Km^R^) gene were cloned into pBR322 to construct the template plasmid. PCR products containing the 5’-end of *polA* and the Km^R^ gene were generated using the primers with long extensions homologous to the upstream *galK*: 1–9 aa region and the downstream *galK*: 21 aa–stop codon region. The PCR products were inserted into the p3478 chromosome, and novel junctions were created between the insert and the neighboring upstream region, as well as between the insert and the downstream region. We randomly isolated two Km^R^ colonies and verified the successful recombination at both of the upstream and the downstream junctions by PCR using locus-specific primers and Sanger sequencing (Fig. 14). As shown in Fig. 14A, the upstream junction was detected using the primer pair galT81aa and kt, both of which were adjacent to the upstream homology. The downstream junction was detected using the primer pair k3 and galK111aa. We used the primer pair galT81aa and galM48aa to further confirm the insertion. If the fragment was successfully inserted, PCR products of 4,524 bp would be observed. Otherwise, if the chromosome remained undisrupted, the PCR products would be 2,094 bp. Because the strains with successful insertions will be further used for recombination in the next “plasmid-in” step, we also PCR confirmed the presence of pSIJ8 at the end of this step. This PCR strategy to detect recombination-directed novel junctions by locus-specific primers was used in the remaining “plasmid-in” steps described below. As shown in Fig. 14A, FRT, “G”, and p1 were also inserted at the downstream junction as auxiliary elements. p1 was the priming site on the template plasmid and remained on the chromosome as a scar after “plasmid-out”. More details about p1 can be found in (Datsenko and Wanner 2000).

**Figure 14.**
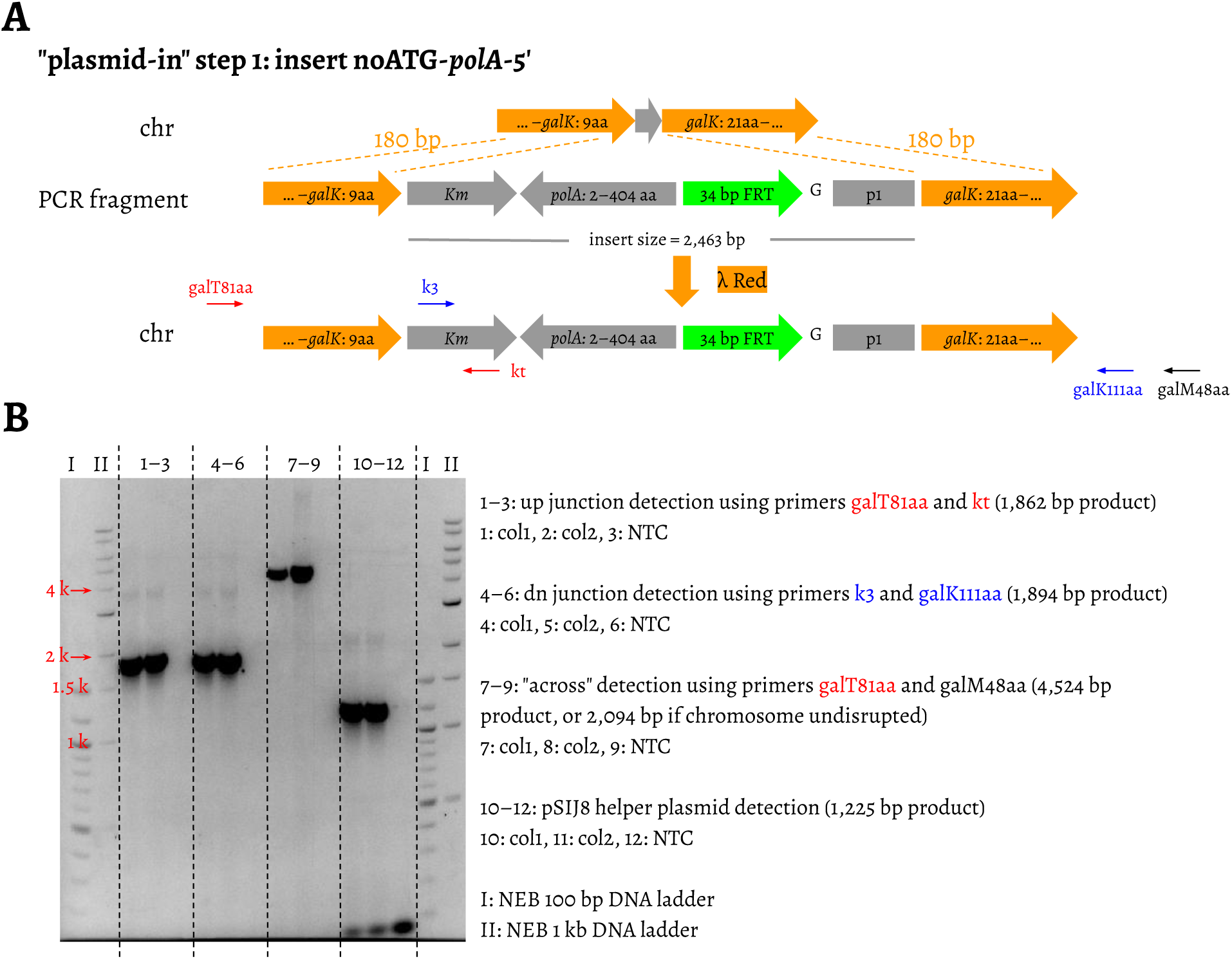

#### “plasmid-in” step 2: insert *polA*3’

In this step, the remaining 3’-end of *polA* and the chloramphenicol resistance (Cm^R^) gene were inserted into the chromosome to replace the Km^R^ gene (Fig. 15). Cm^R^ colonies were isolated and novel junctions created at the recombination upstream and downstream regions were verified by PCR and Sanger sequencing. By the end of this step, *polA* and the downstream auxiliary elements have been fully integrated.

**Figure 15.**
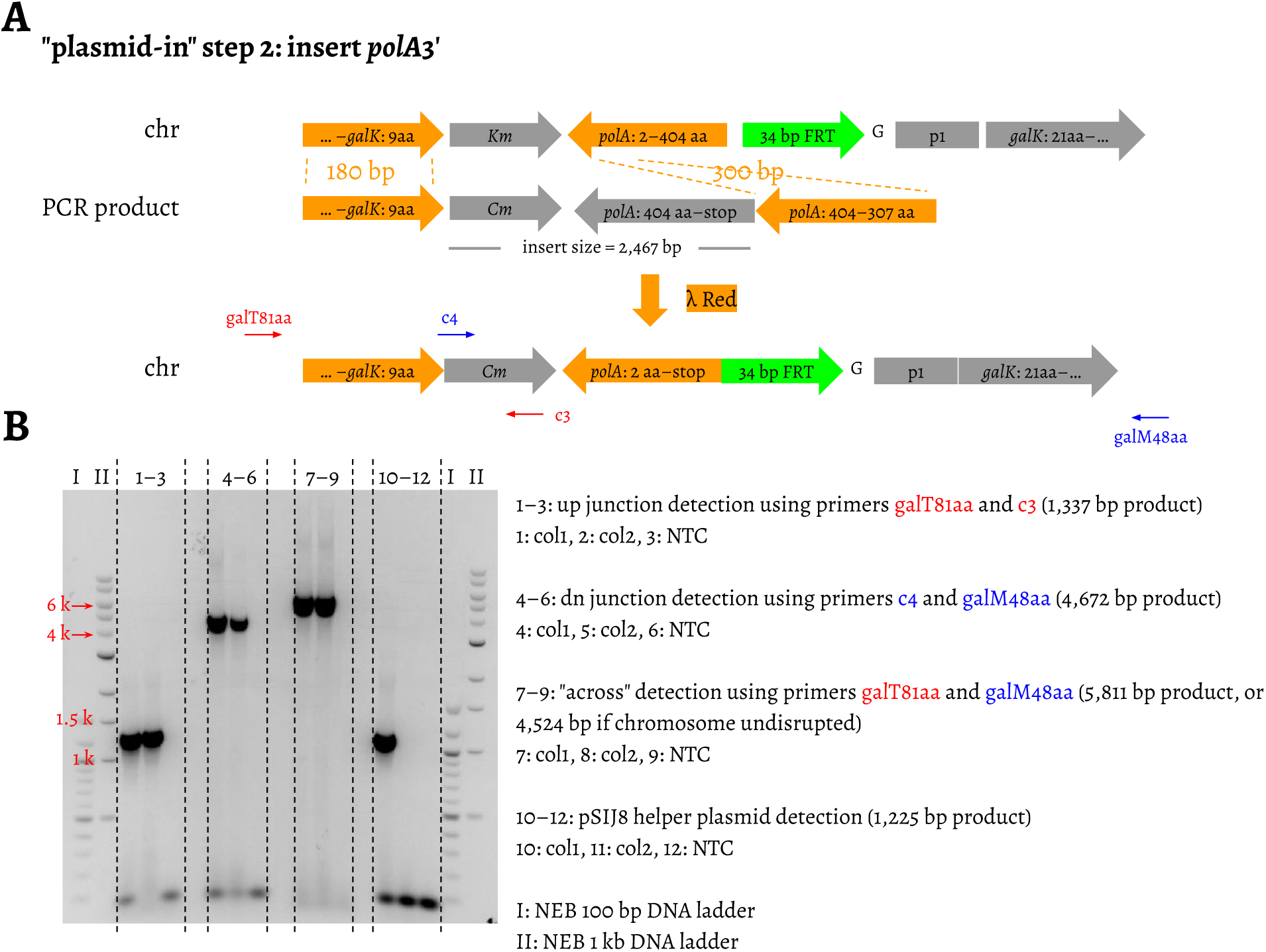

#### “plasmid-in” step 3: insert *ori*

In this step, the pBR322 origin and the Km^R^ gene were inserted into the chromosome to replace the Cm^R^ gene from the last step. We have observed a lot of Km^R^ colonies on the Km agar plate and isolated four colonies to PCR verify the upstream and downstream recombination junctions (Fig. 16). The successful insertion of the pBR322 origin suggested that the *polA* integrated by the end of “plasmid-in” step 2 was inactive.

**Figure 16.**
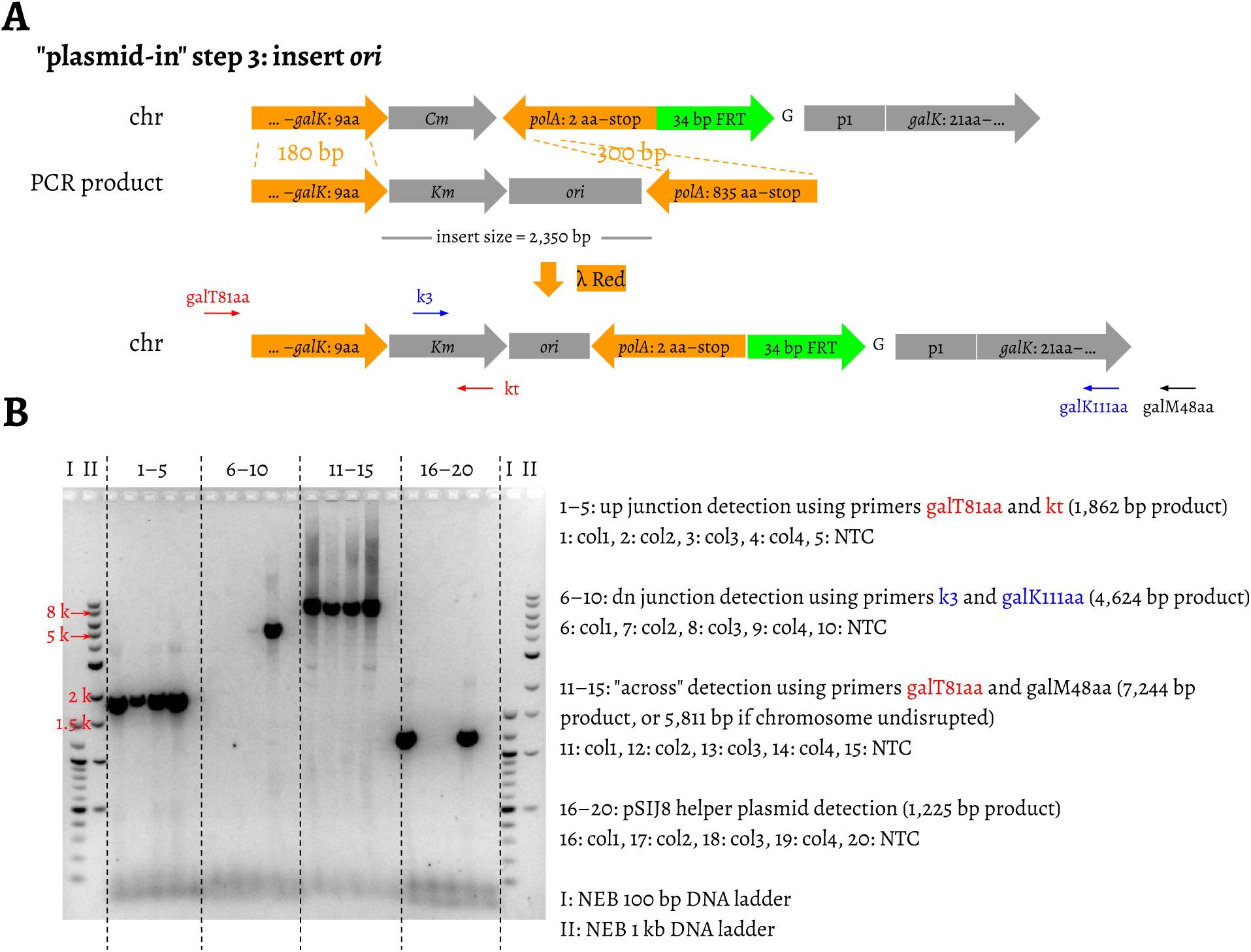

#### “plasmid-in” step 4: insert *rop*-*tetC*3’

After “plasmid-in” steps 4 and 5, by insertion of the *tetC* gene, recombinants would be selected on Gal indicator agar plates supplemented with tetracycline (Tc). The PCR template plasmids in both of these two steps should not bear an intact copy of *tetC* in case that transformants would acquire the template plasmids, specifically residual plasmid molecules surviving the DpnI digestion, and grow as background Tc resistance (Tc^R^) colonies. We split *tetC* into two parts—the 5’-end part *tetC*5’ and the 3’-end part *tetC*3’—at the EagI restriction enzyme site, and put them on the template plasmids of steps 4 and 5 separately. Neither of these two parts could confer a Tc^R^ phenotype alone (Pruss and Drlica 1986). In “plasmid-in” step 4, *rop* and *tetC*3’ together with the Cm^R^ gene were inserted into the chromosome to replace the Km^R^ gene (Fig. 17). The insertion was verified by PCR and Sanger sequencing.

**Figure 17.**
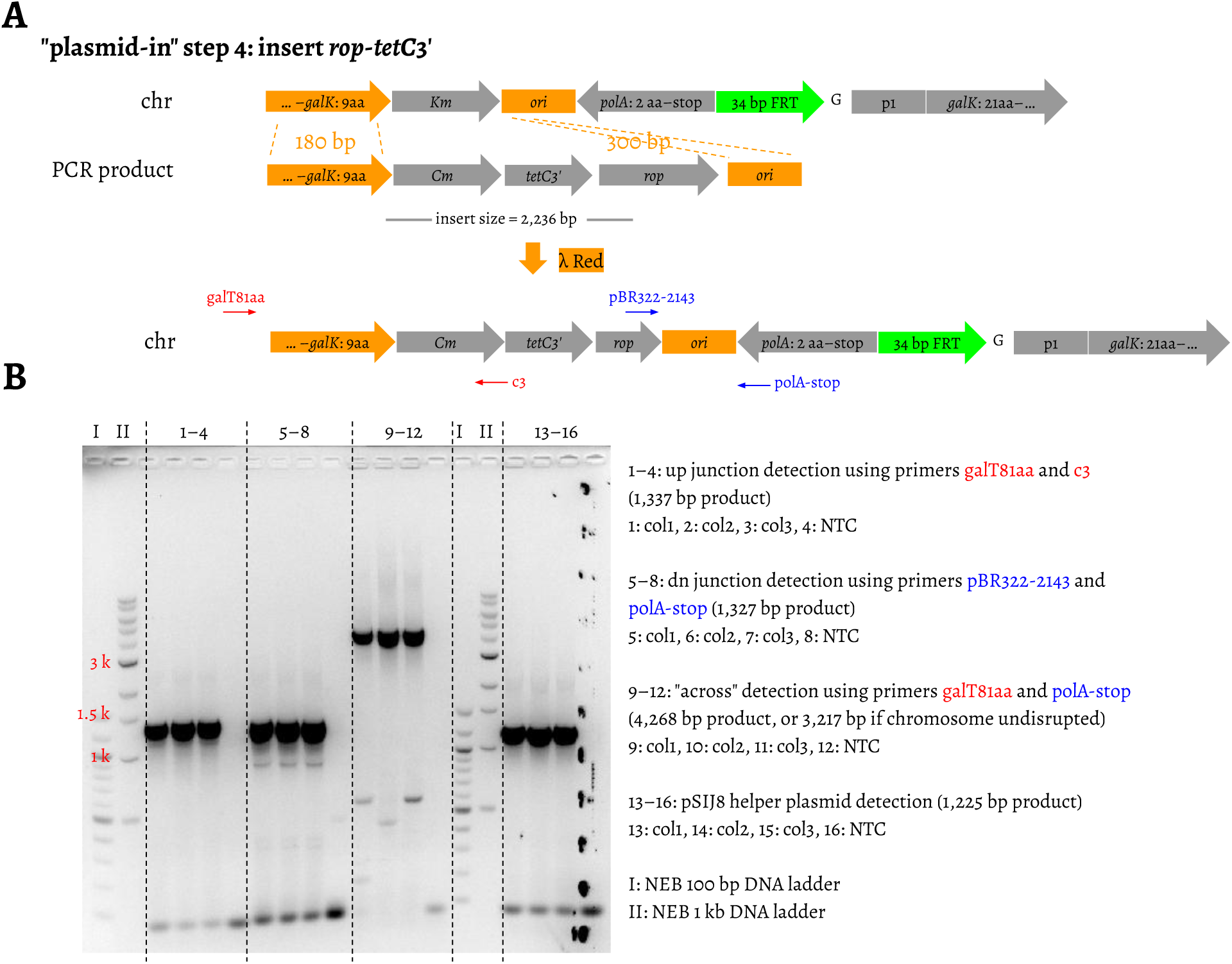

#### “plasmid-in” step 5: insert *barcode*-*tetC*5’

In “plasmid-in” step 5, the goal was to insert *tetC*5’ as well as the chromosomal and plasmid barcodes (Fig. 5A). We cloned the *tetC*5’ fragment into the plasmid carrying *oriRγ* to make the template plasmid. *oriRγ* is a conditional replicative origin, which requires *pir+* for its maintenance, and our template plasmid was propagated in *E. coli* BW25141 (Datsenko and Wanner 2000). p3478 does not carry *pir+*, which ensures that the *oriRγ* template plasmid molecules, if survived DpnI digestion and then electroporated into p3478, would not replicate extrachromosomally. The PCR products were generated from the template plasmid using primers with homology extensions (Fig. 5A). The upstream primer carried the chromosomal and plasmid barcodes, as well as FRT sequence and the auxiliary nucleotides “C” ahead of FRT and “GC” following FRT (Fig. 5A).

After performing “plasmid-in” step 5, we randomly isolated three Tc^R^ colonies—col1, col2, and col3—into LB broth supplemented with Tc, and the cell cultures were denoted as the FLP uninduced cultures. As shown in the lanes marked “FLP uninduced” in Fig. 5C, the novel junctions created by recombination can be detected at both upstream and downstream regions. We also used the primer pair galT81aa and galM48aa, to further confirm the PCR products integration. galT81aa and galM48aa targeted the chromosome and covered across the upstream junction, the whole insert, the downstream junction, and the PCR products should be 8,465 bp. The PCR products using the FLP uninduced col1 culture as the template were subjected to Sanger sequencing to unveil the chromosomal and plasmid barcode combination (Fig.6A).

### Induce FLP expression for plasmid excision

Three colonies—col1, col2, and col3—from “plasmid-in” step 5 were isolated and transferred into LB broth supplemented with 25 μg/mL thymine, 50 μg/mL Ap, and 5 μg/mL Tc, which was used as the FLP uninduced culture. The FLP uninduced culture was further inoculated into the same LB broth, and 50 mM rhamnose were added as the FLP inducer. The cell culture was allowed growth at 30 °C to the late exponential phase, and then used as the FLP induced culture.

### Supplement thiamine and casamino acids

p3478, p3478 transformed with pSIJ8, the resulting strains of “plasmid-in” steps 1–4, and the FLP uninduced and the FLP induced col1, col2, col3 from “plasmid-in” step 5 and “plasmid-out” step 6 were streaked on M63 agar plates supplemented with 25 μg/mL thymine, 0.2% galactose, and the same M63 agar plates with additional supplements of 0.5 μg/mL thiamine hydrochloride, and CAA at the following concentrations: 0.005%, 0.01%, 0.05%, and 0.2%. These plates were incubated at 37 °C.

### Cure pSIJ8 and enrich the recircularized plasmid

The FLP induced col1, col2, and col3 from “plasmid-in” step 5 were inoculated into M63 broth supplemented with 25 μg/mL thymine, 0.2% galactose, 0.5 μg/mL thiamine hydrochloride, 0.05% CAA, 10 μg/mL Tc, 0.2 M NaCl, 1× trace elements A (Corning cat. 25-021-CI), and 1× trace elements B (Corning cat. 25-022-CI). NaCl was added to increase the osmolarity in M63 broth. We found that the trace elements A and B helped improve growth in M63 broth. The cell cultures were grown at 37 °C to late exponential phase, and another round of passaging in the same M63 broth was performed. The plasmids were minipreped, which contained two types of plasmids—pSIJ8 and the recircularized plasmids. pSIJ8 is 9,647 bp, and the recircularized plasmid is 6,329 bp. Both of the two plasmids have only one HindIII restriction site. The minipreped plasmids were cut by HindIII (NEB cat. R3104S), and gel electrophoresed.

### Pilot population barcoding run and ampliconic library preparation

Ten Gal indicator plates (25 μg/mL thymine, 5 μg/mL Tc) after “plasmid-in” step 5 were washed, pooled, and grown at 30 °C to the stationary phase in 70 mL LB supplemented with 25 μg/mL thymine, 50 μg/mL Ap, and 5 μg/mL Tc. This served as the FLP uninduced culture. From the FLP uninduced culture, 15 mL was concentrated and resuspended in 30 mL of the same fresh LB media, with rhamnose added at a final concentration of 50 mM, and grown at 30 °C to the stationary phase. This served as the FLP induced culture. The FLP induced culture was passaged into the M63 and LB line by 1:100 dilution. In the M63 line, 0.3 mL of the FLP induced culture was washed once using the M63 buffer and inoculated into 30 mL M63 broth supplemented with 25 μg/mL thymine, 0.2% galactose, 0.5 μg/mL thiamine hydrochloride, 0.05% CAA, and 10 μg/mL Tc, and were grown at 37 °C to the stationary phase. 0.3 mL of the cell culture was then passaged to 30 mL of the same fresh M63 broth, but with a higher concentration of Tc at 30 μg/mL. In the LB line, 0.3 mL of the FLP induced culture was passaged into 30 mL LB broth supplemented with 25 μg/mL thymine and 10 μg/mL Tc, and was grown at 37 °C to the stationary phase. The cell culture was serially passaged into higher Tc concentrations—30, 50, 100 μg/mL. During each passage, 0.3 mL was added to 30 mL fresh LB media.

Genomic DNA was extracted (Qiagen cat. 51304) from the FLP uninduced and induced cultures, the M63 line sample with 10 μg/mL Tc, and the LB line sample with 100 μg/mL Tc. The genomic DNA was amplified using a two-step PCR process modified from that described in (Levy et al. 2015). In the first step, a 3-cycle PCR was performed using the genomic DNA and Platinum SuperFi II. For the FLP induced sample, both the chromosomal and the plasmid barcodes remained on the chromosome. In the 3-cycle PCR, the oligos *1F-chr* and *1R-chr-uninduced* were used as the forward and the reverse primers, respectively (Table S2). For the remaining samples, the plasmid was excised from the chromosome. In the 3-cycle PCR, for the chromosome, the oligos *1F-chr* and *1R-chr-FLP-induced* were used as the forward and the reverse primers, respectively (Table S2). In the 3-cycle PCR, for the plasmid, the oligos *1F-plasmid* and *1R-plasmid* were used as the forward and the reverse primers, respectively (Table S2).

The Ns in the primers are random nucleotides and are used to remove the bias in DNA molecule counting—referred to as PCR jack-potting. The 3-cycle PCR products were purified using 1.8× beads, and resuspended in H2O. The purified 3-cycle PCR products were qPCR quantified using Platinum SuperFi II on a CFX96 system (BioRad). The same amount of the seven purified 3-cycle PCR products were further amplified using Platinum SuperFi II and the index primers from Nextera XT Index Kit v2 (Illumina cat. 15052163). The PCR products were purified using 1.8× beads, and resuspended in 10 mM Tris-HCl, which were used as the completed libraries. The FLP uninduced chromosome library was denoted as D2_P1. The FLP induced chromosomal and plasmid libraries were denoted as D2_P2 and D2_P3 respectively. For the M63 line sample at Tc 10 μg/mL, the chromosomal and plasmid libraries were denoted as D2_P4 and D2_P5 respectively. For the LB line sample at Tc 100 μg/mL, the chromosomal and plasmid libraries were denoted as D2_P6 and D2_P7 respectively. The seven completed libraries D2_P1–D2_P7 were checked by Bioanalyzer (Agilent), quantified using qPCR, made an approximately equimolar pool, and sequenced in a Nextseq 550 75 × 75 mid output run with 30% phiX spike-in.

### Nextseq 550 data analysis

Reads with base quality below Q20 were removed. We then filtered out the reads if they did not match the reads structures as shown in Fig. 9 with no mismatch allowed. The PCR jack-potting barcodes on the read1 and read2 sides were referred to as JP1 and JP2 respectively. The JP1 and JP2 barcodes were combined to form JP1-JP2 families. For any unique JP1-JP2 family, the family size was defined as the appearance of the JP1-JP2 combination. In the manuscript, we defined lineage size as the number of DNA molecules in a lineage. Lineage size was obtained as described below. For D2_P1, the filtered reads were grouped by chromosome-plasmid barcode combinations, and then grouped by JP1-JP2 combination. We next applied a JP1-JP2 family threshold of 2, and removed JP1-JP2 families with a family size of 1. JP1-JP2 families were then merged to form JP1-JP2 consensus sequences. For any chromosome-plasmid barcode combination lineage, the lineage size was the number of JP1-JP2 consensus sequences in that lineage. It should be noted that in D2_P1, we only kept the unique chromosome-plasmid barcode combinations. For D2_P2–D2_P7, the reads were grouped by either chromosome or plasmid barcodes, with no JP1-JP2 family size threshold applied. The lineage size of each chromosome or plasmid barcode was obtained in the same way as D2_P1 by counting JP1-JP2 families. The jupyter notebook “Barcode_deconvolution.ipynb” to deconvolute the UMI-tagged reads and assess the barcode complexity can be found at https://github.com/nekrut/plasmid.

## Data availability

This project has been deposited at Short Read Archive (SRA) at the National Center for Biotechnology Information (NCBI) under the BioProject accession PRJNA794739. SRA records are under the same D2_P1–D2_P7 denotations as in the manuscript.

## Supplementary files

Supplementary files include the following:

- Table S1 (plasmids used in this study, xlsx format),
- Table S2 (primers used in this study, xlsx format),
- Table S3 (lineages size recovered in the pilot run, xlsx format),
- maps of the *galK* fusion plasmids to explore the *galK* integration site (SnapGene Viewer dna format),
- maps of the plasmids to test if *galK* activates *polA* expression (SnapGene Viewer dna format),
- PCR template plasmid maps, the p3478 chromosomal structures ranging from *galT* Ala81 to *galM* Ser48 of the resulting strains from “plasmid-in” step 1–5, and the chromosomal and the recircularized plasmid structures after “plasmid-out” step 6 (SnapGene Viewer dna format),
- the UMI-tagged ampliconic libraries’ structures (SnapGene Viewer dna format).

## Acknowledgements

The authors are grateful to Dr. Paul Babitzke, Dr. Kenneth Keiler, Dr. Lu Bai, and Hengye Chen from the BMB department at Penn State for offering molecular biology advice during siBar development and to Dr. Stéphanie Bedhomme for critical reading of the manuscript. We thank Dr. Craig A. Praul for advising on Nextera library preparation and Penn State Genomics Core Facility - University Park, PA for providing Sanger sequencing and Nextseq sequencing services. This study has been funded by the funds provided by the Eberly College of Science at the Pennsylvania State University.

## References

Baba T, Ara T, Hasegawa M, Takai Y, Okumura Y, Baba M, Datsenko KA, Tomita M, Wanner BL, Mori H. 2006. Construction of Escherichia coli K-12 in-frame, single-gene knockout mutants: the Keio collection. Mol. Syst. Biol. 2:2006.0008.

Bedhomme S, Perez Pantoja D, Bravo IG. 2017. Plasmid and clonal interference during post horizontal gene transfer evolution. Mol. Ecol. 26:1832–1847.

Broom M, Krivan V. 2018. Biology and evolutionary games.

Csiszovszki Z, Krishna S, Orosz L, Adhya S, Semsey S. 2011. Structure and function of the D-galactose network in enterobacteria. MBio 2:e00053–11.

Datsenko KA, Wanner BL. 2000. One-step inactivation of chromosomal genes in Escherichia coli K-12 using PCR products. Proc. Natl. Acad. Sci. U. S. A. 97:6640–6645.

DeVito JA. 2008. Recombineering with tolC as a selectable/counter-selectable marker: remodeling the rRNA operons of Escherichia coli. Nucleic Acids Res. 36:e4.

Farrell MJ, Finkel SE. 2003. The Growth Advantage in Stationary-Phase PhenotypeConferred by rpoS Mutations Is Dependent on the pH andNutrientEnvironment. Journal of Bacteriology [Internet] 185:7044–7052. Available from: http://dx.doi.org/10.1128/jb.185.24.7044-7052.2003

Garoña A, Hülter NF, Romero Picazo D, Dagan T. 2021. Segregational drift constrains the evolutionary rate of prokaryotic plasmids. Mol. Biol. Evol. [Internet]. Available from: http://dx.doi.org/10.1093/molbev/msab283

Gross J, Gross M. 1969. Genetic Analysis of an E. coli Strain with a Mutation affecting DNA Polymerase. Nature [Internet] 224:1166–1168. Available from: http://dx.doi.org/10.1038/2241166a0

Gutterson NI, Koshland DE Jr. 1983. Replacement and amplification of bacterial genes with sequences altered in vitro. Proc. Natl. Acad. Sci. U. S. A. 80:4894–4898.

Hartley A, Glynn SE, Barynin V, Baker PJ, Sedelnikova SE, Verhees C, de Geus D, van der Oost J, Timson DJ, Reece RJ, et al. 2004. Substrate specificity and mechanism from the structure of Pyrococcus furiosus galactokinase. J. Mol. Biol. 337:387–398.

Ilhan J, Kupczok A, Woehle C, Wein T, Hülter NF, Rosenstiel P, Landan G, Mizrahi I, Dagan T. 2019. Segregational Drift and the Interplay between Plasmid Copy Number and Evolvability. Mol. Biol. Evol. 36:472–486.

Jahn M, Vorpahl C, Hübschmann T, Harms H, Müller S. 2016. Copy number variability of expression plasmids determined by cell sorting and Droplet Digital PCR. Microb. Cell Fact. 15:211.

Jensen SI, Lennen RM, Herrgård MJ, Nielsen AT. 2015. Seven gene deletions in seven days: Fast generation of Escherichia coli strains tolerant to acetate and osmotic stress. Sci. Rep. 5:17874.

Jensen SI, Nielsen AT. 2018. Multiplex Genome Editing in Escherichia coli. Methods Mol. Biol. 1671:119–129.

Joyce CM, Kelley WS, Grindley ND. 1982. Nucleotide sequence of the Escherichia coli polA gene and primary structure of DNA polymerase I. J. Biol. Chem. 257:1958–1964.

Kingsbury DT, Helinski DR. 1970. DNA polymerase as a requirement for the maintenance of the bacterial plasmid colicinogenic factor E1. Biochemical and Biophysical Research Communications [Internet] 41:1538–1544. Available from: http://dx.doi.org/10.1016/0006-291x(70)90562-0

Kuhlman TE, Cox EC. 2010. Site-specific chromosomal integration of large synthetic constructs. Nucleic Acids Res. 38:e92.

Lee EC, Yu D, Martinez de Velasco J, Tessarollo L, Swing DA, Court DL, Jenkins NA, Copeland NG. 2001. A highly efficient Escherichia coli-based chromosome engineering system adapted for recombinogenic targeting and subcloning of BAC DNA. Genomics 73:56–65.

Levy SF, Blundell JR, Venkataram S, Petrov DA, Fisher DS, Sherlock G. 2015. Quantitative evolutionary dynamics using high-resolution lineage tracking. Nature 519:181–186.

Li X-T, Thomason LC, Sawitzke JA, Costantino N, Court DL. 2013. Positive and negative selection using the tetA-sacB cassette: recombineering and P1 transduction in Escherichia coli. Nucleic Acids Res. 41:e204.

de Lucia P, Cairns J. 1969. Isolation of an E. coli Strain with a Mutation affecting DNA Polymerase. Nature [Internet] 224:1164–1166. Available from: http://dx.doi.org/10.1038/2241164a0

Maresca M, Erler A, Fu J, Friedrich A, Zhang Y, Francis Stewart A. 2010. Single-stranded heteroduplex intermediates in λ Red homologous recombination. BMC Molecular Biology [Internet] 11:54. Available from: http://dx.doi.org/10.1186/1471-2199-11-54

Mei H, Arbeithuber B, Cremona MA, DeGiorgio M, Nekrutenko A. 2019. A High-Resolution View of Adaptive Event Dynamics in a Plasmid. Genome Biology and Evolution [Internet] 11:3022–3034. Available from: http://dx.doi.org/10.1093/gbe/evz197

Mosberg JA, Lajoie MJ, Church GM. 2010. Lambda red recombineering in Escherichia coli occurs through a fully single-stranded intermediate. Genetics 186:791–799.

Paulsson J. 2002. Multileveled selection on plasmid replication. Genetics 161:1373–1384.

Pruss GJ, Drlica K. 1986. Topoisomerase I mutants: the gene on pBR322 that encodes resistance to tetracycline affects plasmid DNA supercoiling. Proc. Natl. Acad. Sci. U. S. A. 83:8952–8956.

Rodríguez-Beltrán J, DelaFuente J, León-Sampedro R, MacLean RC, San Millán Á. 2021. Beyond horizontal gene transfer: the role of plasmids in bacterial evolution. Nat. Rev. Microbiol. 19:347–359.

Rodriguez-Beltran J, Hernandez-Beltran JCR, DelaFuente J, Escudero JA, Fuentes-Hernandez A, MacLean RC, Peña-Miller R, San Millan A. 2018. Multicopy plasmids allow bacteria to escape from fitness trade-offs during evolutionary innovation. Nat Ecol Evol 2:873–881.

Saarilahti HT, Tapio Palva E. 1985. In vivo transfer of chromosomal mutations onto multicopy plasmids utilizing polA strains: Cloning of an ompR 2 mutation in Escherichia coli K-12. FEMS Microbiology Letters [Internet] 26:27–33. Available from: http://dx.doi.org/10.1111/j.1574-6968.1985.tb01560.x

Sezonov G, Joseleau-Petit D, D’Ari R. 2007. Escherichia coli physiology in Luria-Bertani broth. J. Bacteriol. 189:8746–8749.

Smith JM. 1982. Evolution and the Theory of Games. Cambridge University Press

Stüber D, Bujard H. 1981. Organization of transcriptional signals in plasmids pBR322 and pACYC184. Proc. Natl. Acad. Sci. U. S. A. 78:167–171.

Turan S, Bode J. 2011. Site-specific recombinases: from tag-and-target- to tag-and-exchange-based genomic modifications. FASEB J. 25:4088–4107.

Wang H, Bian X, Xia L, Ding X, Müller R, Zhang Y, Fu J, Stewart AF. 2014. Improved seamless mutagenesis by recombineering using ccdB for counterselection. Nucleic Acids Res. 42:e37.

Warming S, Costantino N, Court DL, Jenkins NA, Copeland NG. 2005. Simple and highly efficient BAC recombineering using galK selection. Nucleic Acids Res. 33:e36.

Wong Ng J, Chatenay D, Robert J, Poirier MG. 2010. Plasmid copy number noise in monoclonal populations of bacteria. Phys. Rev. E Stat. Nonlin. Soft Matter Phys. 81:011909.

Wong QNY, Ng VCW, Lin MCM, Kung H-F, Chan D, Huang J-D. 2005. Efficient and seamless DNA recombineering using a thymidylate synthase A selection system in Escherichia coli. Nucleic Acids Res. 33:e59.

Zhang Y, Muyrers JPP, Rientjes J, Stewart AF. 2003. Phage annealing proteins promote oligonucleotide-directed mutagenesis in Escherichia coli and mouse ES cells. BMC Mol. Biol. 4:1.

